# *In vivo* genetic screen identifies a SLC5A3-dependent myo-inositol auxotrophy in acute myeloid leukemia

**DOI:** 10.1101/2020.12.22.424018

**Authors:** Yiliang Wei, Shruti V. Iyer, Ana S. H. Costa, Zhaolin Yang, Melissa Kramer, Emmalee R. Adelman, Olaf Klingbeil, Osama E. Demerdash, Sofya Polyanskaya, Kenneth Chang, Sara Goodwin, Emily Hodges, W. Richard McCombie, Maria E. Figueroa, Christopher R. Vakoc

## Abstract

An enhanced requirement for extracellular nutrients is a hallmark property of cancer cells. Here, we optimized an *in vivo* genetic screening strategy for evaluating dependencies in acute myeloid leukemia (AML), which led to the identification of the myo-inositol transporter SLC5A3 as a unique vulnerability in this disease. In accord with this transport function, we demonstrate that the SLC5A3 dependency reflects a myo-inositol auxotrophy in AML. Importantly, the commonality among SLC5A3-dependent AML lines is the transcriptional silencing of *ISYNA1*, which encodes the rate limiting enzyme for myoinositol biosynthesis, inositol-3-phosphate synthase 1. We used gain- and loss-of-function experiments to demonstrate a synthetic lethal genetic interaction between *ISYNA1* and *SLC5A3* in AML, which function redundantly to sustain intracellular myo-inositol. Transcriptional silencing and DNA hypermethylation of *ISYNA1* occur in a recurrent manner in human AML patient samples, in association with the presence of *IDH1/IDH2* and *CEBPA* mutations. Collectively, our findings reveal myo-inositol auxotrophy as a novel form of metabolic dysregulation in AML, which is caused by the aberrant silencing of a biosynthetic enzyme.

**Statement of significance:** Here, we show how epigenetic silencing can provoke a nutrient dependency in AML by exploiting a synthetic lethality relationship between biosynthesis and transport of myo-inositol. Blocking the function of this solute carrier may have therapeutic potential in an epigenetically-defined subset of AML.

## Introduction

A consistent property of cancer cells is metabolic dysfunction, which can occur to support the unique energetic and biosynthetic requirements of a rapidly growing tumor (DeBerardinis & Chandel 2016). One manifestation of this dysfunction is that tumor cells become more dependent on extracellular nutrient uptake for their growth and viability than normal tissues, a process referred to as auxotrophy (Garcia-Bermudez et al. 2020). In many tumors, auxotrophies are driven by an elevated anabolic demand for metabolite building blocks, as is the case for serine (Possemato et al. 2011), glutamine (Wise et al. 2008), and cysteine (Cramer et al. 2017). However, it has been shown that cancer cells can develop nutrient dependencies through an acquired defect in *de novo* metabolite biosynthesis pathways. One of the clearest examples of this is the dependence of lymphoid cancers on extracellular asparagine, which occurs because *ASNS*, encoding the rate limiting enzyme in asparagine biosynthesis, becomes epigenetically silenced in this malignancy (Peng et al. 2001). This nutrient dependency can be exploited by infusion of recombinant asparaginase, which is an approved chemotherapeutic in acute lymphoblastic leukemia that depletes asparagine from the plasma (Jaffe et al. 1971). Arginine and cholesterol auxotrophies have also been described in cancer and attributed to diminished expression of key biosynthetic enzymes (Nicholson et al. 2009, Garcia-Bermudez et al. 2019). While these examples point to the therapeutic significance of auxotrophy in human cancer, few systematic surveys of nutrient dependencies have been performed in cancer to date.

Solute carriers (SLC) are an important class of transporter proteins that regulate nutrient utilization in normal and neoplastic contexts. The human genome encodes over 400 SLC proteins organized into 65 subfamilies, which regulate intracellular levels of amino acids, sugars, ions, lipids, neurotransmitters, and drugs (Nakanishi & Tamai 2011). SLCs contain transmembrane alpha helices that form a substrate binding channel, which often use ion gradients to co-transport substrates into cells (Bai et al. 2017). Owing to their specificity for transporting specific substrates, genetic targeting of SLCs provides a powerful tool for manipulation of intracellular metabolite levels and for exploration of nutrient dependencies (Ganapathy et al. 2009, Bacci et al. 2019). Numerous SLC genes are mutated in human genetic disorders and an emerging body of evidence suggests a role for SLC proteins in the pathogenesis of human cancer (El-Gebali et al. 2013, Lin et al. 2015, Zhang et al. 2019). Moreover, the substrate binding cavity of SLC proteins can be selectively inhibited with small molecules, which has fueled an interest in this class of proteins as therapeutic targets (Nyquist et al. 2017, Zhang et al. 2019, Hu et al. 2020).

Myo-inositol is an abundant carbocyclic sugar alcohol in eukaryotic cells (Holub 1986, Croze & Soulage 2013). One of the most well-characterized functions of myo-inositol is as a head group component of lipid phosphatidylinositols (PIs), which are important signaling molecules that function downstream of growth factor signaling. In this context, the phosphorylation state of myo-inositol within PI is highly regulated by kinases (e.g. PI3K) and phosphatases (e.g. PTEN) to control the AKT-mTOR signaling axis (Martelli et al. 2010). In addition, phospholipase C can release inositol triphosphates from PIs, which function as a second messenger that promotes the intracellular release of calcium from the endoplasmic reticulum (Berridge & Irvine 1989, Bansal & Majerus 1990). Myo-inositol is also an osmolyte, whose intracellular concentrations can be regulated to allow cells to survive in hypertonic environments (Nakanishi et al. 1988, Kitamura et al. 1997).

Intracellular myo-inositol can be derived from three different sources: (1) *de novo* biosynthesis from glucose (Eisenberg & Bolden 1963, Hauser & Finelli 1963); (2) regeneration from inositol phosphates via the action of phosphatases (Irvine & Schell 2001); or (3) uptake from the extracellular environment by inositol transporters (Schneider 2015). For *de novo* biosynthesis, glucose is first phosphorylated by hexokinase and then converted to myo-inositol-1-phosphate by the enzyme inositol-3-phosphate synthase 1 (ISYNA1), and finally dephosphorylated by inositol monophosphatase to form free myo-inositol. Importantly, ISYNA1 catalyzes the rate-limiting step in myo-inositol biosynthesis (Stein & Geige 2002). For extracellular uptake, there are three known myo-inositol SLC transporters in human, SLC5A3 (Kwon et al. 1992), SLC5A11 (Hitomi & Tsukagoshi 1994), and SLC2A13 (Uldry et al. 2001). Both SLC5A3 and SLC5A11 are sodium ion coupled inositol transporters, whereas SLC2A13 is a proton-coupled inositol transporter (Schneider 2015). Among all three transporters, SLC5A3 is the most widely expressed across different tissues, and has the highest affinity for myo-inositol (Berry et al. 1995, Hager et al. 1995, Roll et al. 2002, Uldry et al. 2001, Coady et al. 2002).

In this study, we set out to identify genetic dependencies needed for the growth of acute myeloid leukemia cells *in vivo.* To this end, we established a robust domain-focused CRISPR screening strategy in an AML xenograft model. This approach identified the myo-inositol transporter SLC5A3 as an *in vivo*-relevant dependency unique to AML. We show that this dependency is caused by aberrant silencing of *ISYNA1*. While loss of *ISYNA1* has no detectable effect on the fitness of AML cells, it causes an enhanced dependency on SLC5A3 and on extracellular myo-inositol. Taken together, our findings reveal myo-inositol auxotrophy as metabolic vulnerability in AML.

## Results

### A domain-focused CRISPR screening method for identifying *in vivo*-relevant AML dependencies

The genetic dependencies of a cancer cell can be markedly altered by *in vitro* versus *in vivo* growth conditions (Rossiter et al. 2020), particularly the requirement for cell surface receptors and metabolic enzymes whose function is influenced by the extracellular microenvironment. This issue provided the rationale for developing a robust screening method for interrogating genetic dependencies of human AML cells grown *in vivo*. For this study, we employed our previously cloned domain-focused sgRNA libraries targeting transcription factors (Lu et al. 2018), chromatin regulators (Brien et al. 2018, Lan et al. 2019), and kinases (Tarumoto et al. 2018). For these *in vivo* screens, we also generated domain-focused sgRNA libraries targeting G-protein coupled receptors, ion channels, solute carriers, glycosyltransferases, proteases, and membrane proteins, which were all cloned into the LRG2.1T vector backbone which contains an optimized sgRNA2.1 scaffold (Grevet et al. 2018). Our screens were performed by lentiviral transduction of each sgRNA library into the AML cell line MOLM-13, engineered to express Cas9 and luciferase. A key optimization step in establishing this protocol was to select for high engraftment efficiency of this MOLM-13 line through one round of transplantation before running the pooled screen (Figure 1A). After transduction with an sgRNA library, MOLM-13 cells were injected via tail vein into 10-20 immune-deficient mice. After expansion of MOLM-13 cells for 12 days *in vivo*, mice were sacrificed and genomic DNA was isolated from bone marrow and spleen tissue, followed by deep sequencing of the PCR-amplified sgRNA cassette (Figure 1A). While establishing this procedure, we found that pooling of bone marrow and spleen samples prior to PCR maximized sgRNA representation and the overall accuracy of the screens. Importantly, the sgRNA read distribution (a reflection of screen quality) and the depletion pattern of control sgRNAs of *in vivo* samples was comparable to the same screen being performed *in vitro* (Figure S1, Supplemental file 1). We have previously characterized AML vulnerabilities *in vitro* using sgRNA libraries targeting kinases (Tarumoto et al. 2018), transcription factors (Lu et al. 2018), and chromatin regulators (Brien et al. 2018, Lan et al. 2019). The *in vivo* CRISPR screening showed highly consistent results with our previous findings (Figure 1B-D). The overall accuracy of our screen was further supported by the *in vivo*-specific depletion of sgRNAs targeting *CD47* (Oldenborg et al. 2000, Blazar et al. 2001, Sick et al. 2012) and *CXCR4* (Burger & Kipps 2006), which are known AML dependencies that encode membrane proteins regulating interactions with the *in vivo* tissue microenvironment (Figure 1E, Figure S7D-F). In contrast to previous *in vivo* RNAi screening in leukemia that shows only ~10% of *in vivo* dependencies overlapping with *in vitro* targets (Meacham et al. 2015), our screening showed largely consistent results between *in vivo* and *in vitro* conditions (Figure 1F, Figure S2, Supplemental file 1). From this screening effort targeting a total of 5,768 genes across these libraries, we identified 239 targets as *in vivo*-relevant dependencies in MOLM-13 cells (Figure 1F, Supplemental file1).

**Figure 1.**
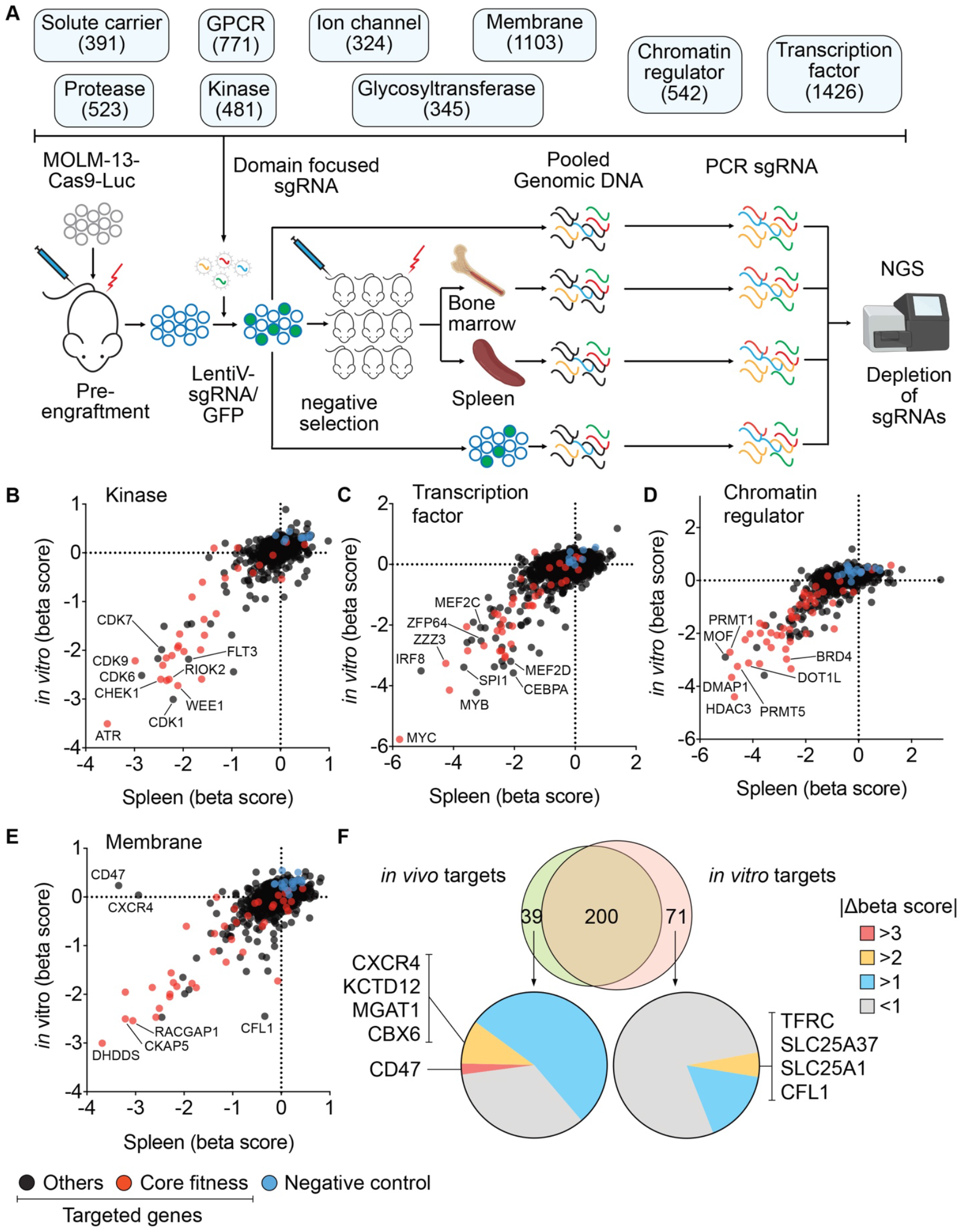
A domain-focused CRISPR screening method for identifying *in vivo*-relevant AML dependencies. (A) Overview of *in vivo* genetic screening procedure using domain-focused sgRNA libraries. GPCR, G-protein coupled receptor; NGS, Next Gen Sequencing. (B-E) Examples of *in vivo* CRISPR screening results with sgRNA libraries targeting kinases (B), transcription factors (C), chromatin regulators (D), and membrane proteins (E). The gene beta score was calculated using MAGeCK (Li et al. 2014). The beta score describes how the gene knockout alters cell fitness: a positive beta score indicates positive selection, and a negative beta score indicates negative selection. The screening results using pooled spleen samples were compared with *in vitro* screening results. Core fitness genes (Hart et al. 2015) are labeled in red, negative control sgRNAs are labeled in blue. (F) Venn diagram comparing targets identified by *in vivo* screening with *in vitro* screening. The *in vivo*-relevant targets were determined using spleen data with *p*>-value <0.05, FDR <0.25, beta score <0. For *in vivo-* and *in vitro*-specific targets, the absolute value of differential beta scores were shown as pie charts. Genes with more than two differential beta scores were listed.

**Figure S1.**
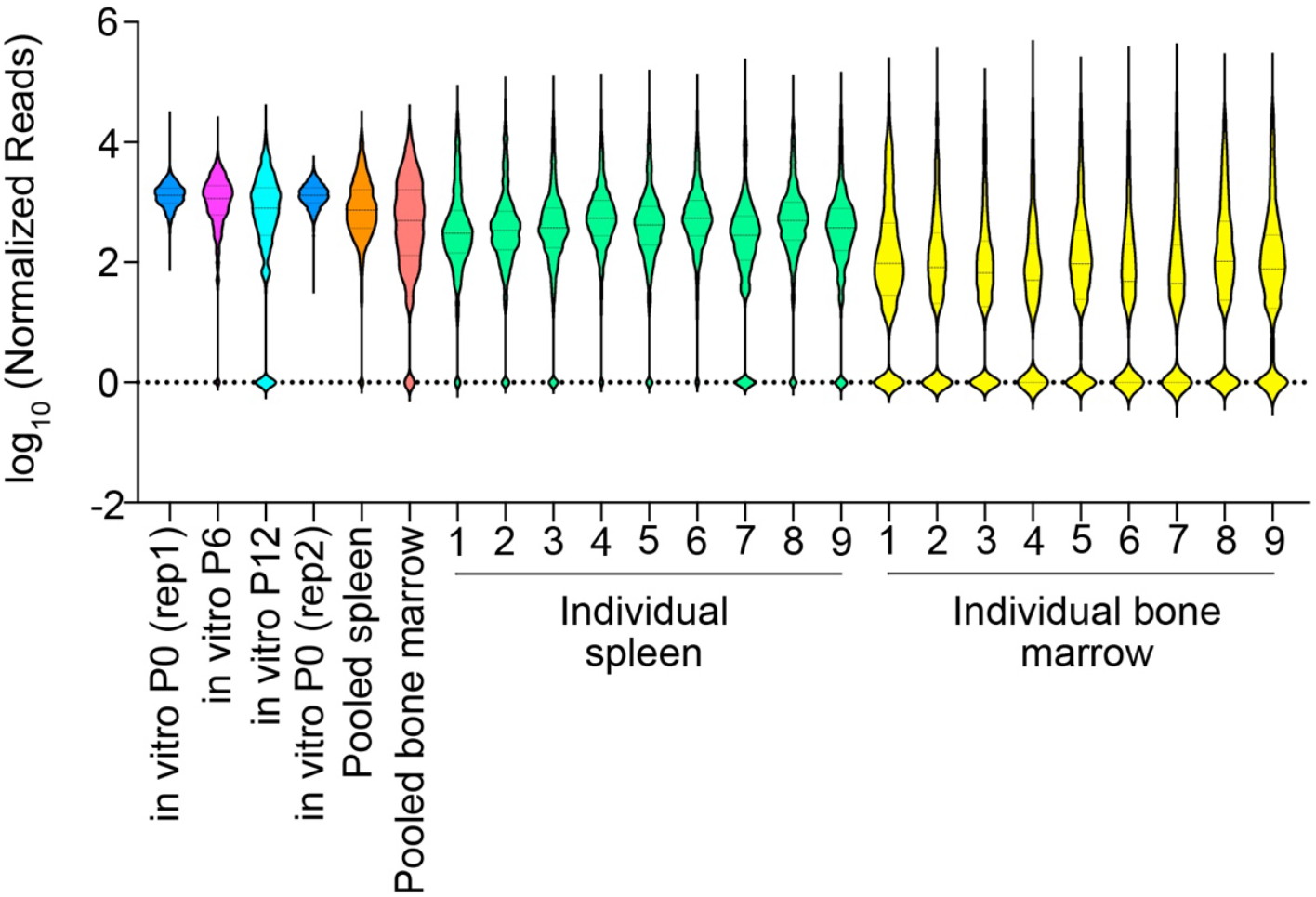
Violin plot of normalized read counts of all sgRNA from the screening samples. P0, passage 0; P6 passage 6; P12, passage 12.

**Figure S2.**
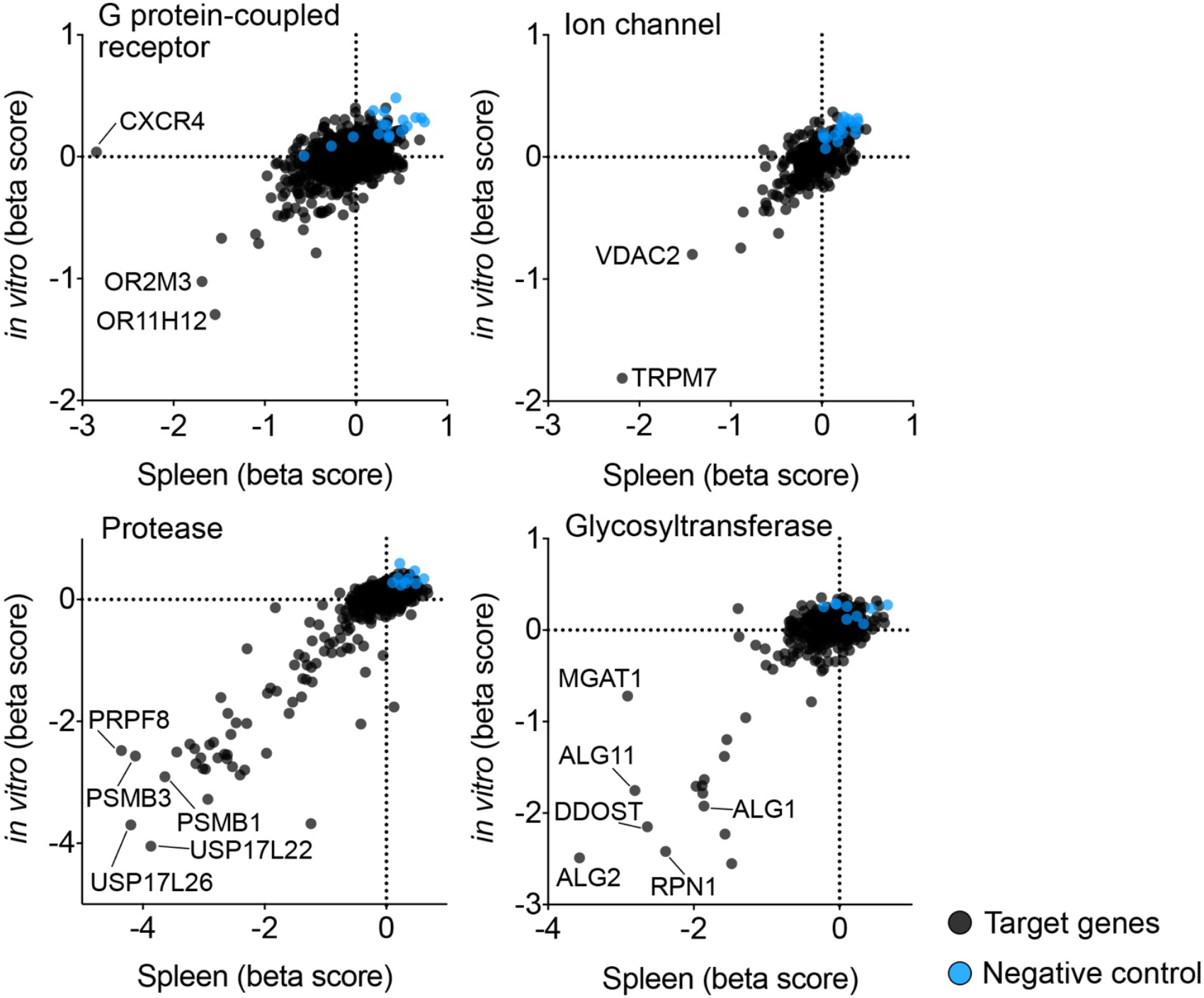
Summary of *in vivo* CRISPR screening results, plotted as beta scores from pooled spleen samples vs. *in vitro* samples.

**Figure S3.**
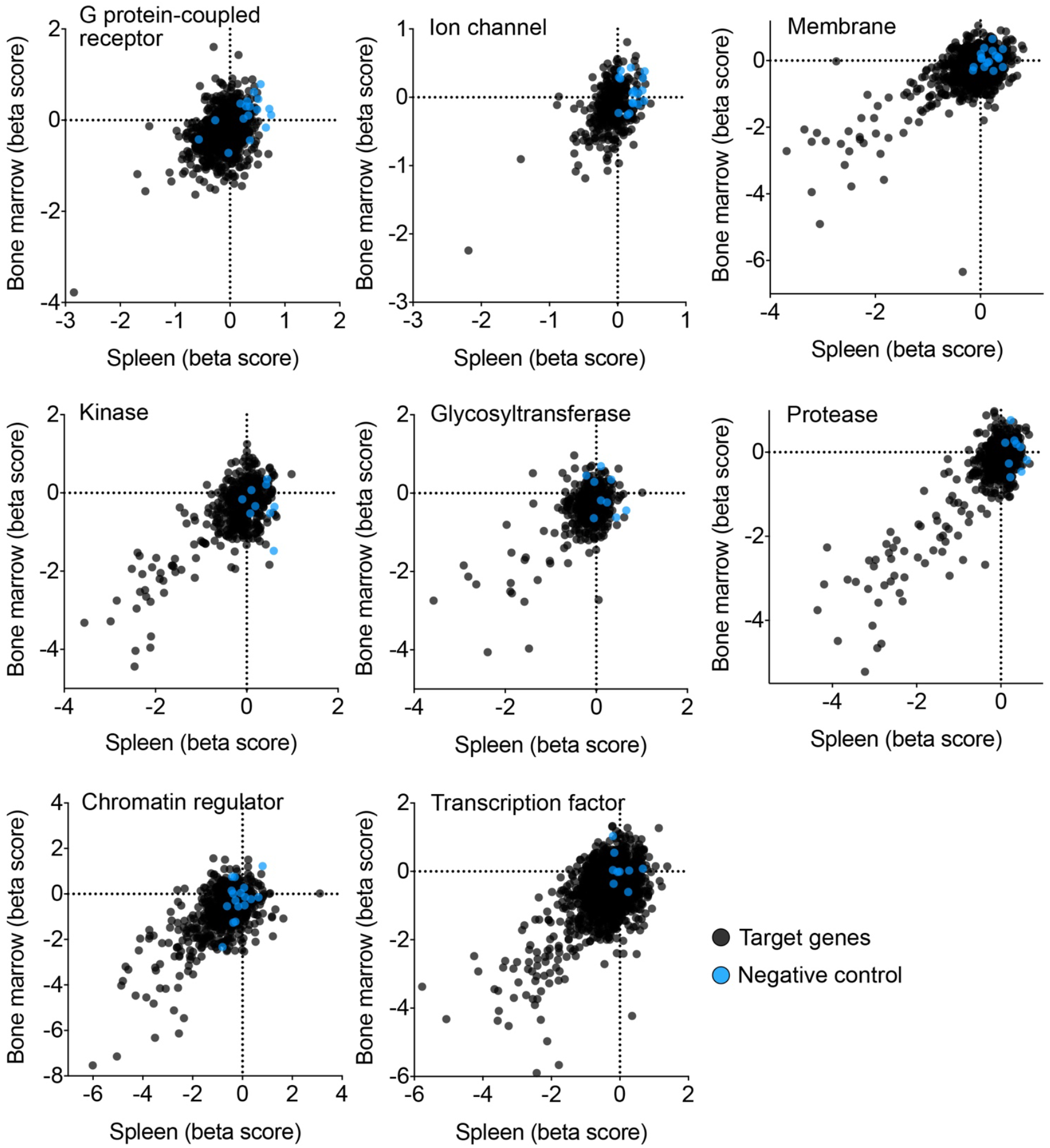
Summary of *in vivo* CRISPR screening results, plotted as beta scores from pooled spleen samples vs. pooled bone marrow samples.

### SLC5A3 dependency is *in vivo*-relevant and AML-specific

We next evaluated how specific the *in vivo*-relevant dependencies were for the leukemia context relative to other cancer types. For this purpose, we turned to Project Achilles, which performed genomewide CRISPR screening for effects on cell fitness in 769 diverse cancer cell lines grown *in vitro* (Meyers et al. 2017, Dempster et al. 2019). Through this analysis, we nominated several myeloid lineage TFs (MYB, CBFB, and SPI1/PU.1) and the myo-inositol transporter protein SLC5A3 as the most leukemia-biased dependencies among our pool of *in vivo*-validated targets (Figure 2A). The AML specificity of SLC5A3 essentiality was distinct from most other SLC genes, which tended to be pan-essential across diverse cancer types (Figure 2C). The *in vivo* screening in MOLM-13 cells and our own *in vitro* SLC screening in 10 diverse cancer cell lines corroborated SLC5A3 as powerful and unique AML dependency under both *in vitro* and *in vivo* conditions (Figure 2B, Figure S4). Unlike the aforementioned myeloid TFs, no prior study has investigated the role for SLC5A3 in AML, which motivated our subsequent evaluation of this target.

**Figure 2.**
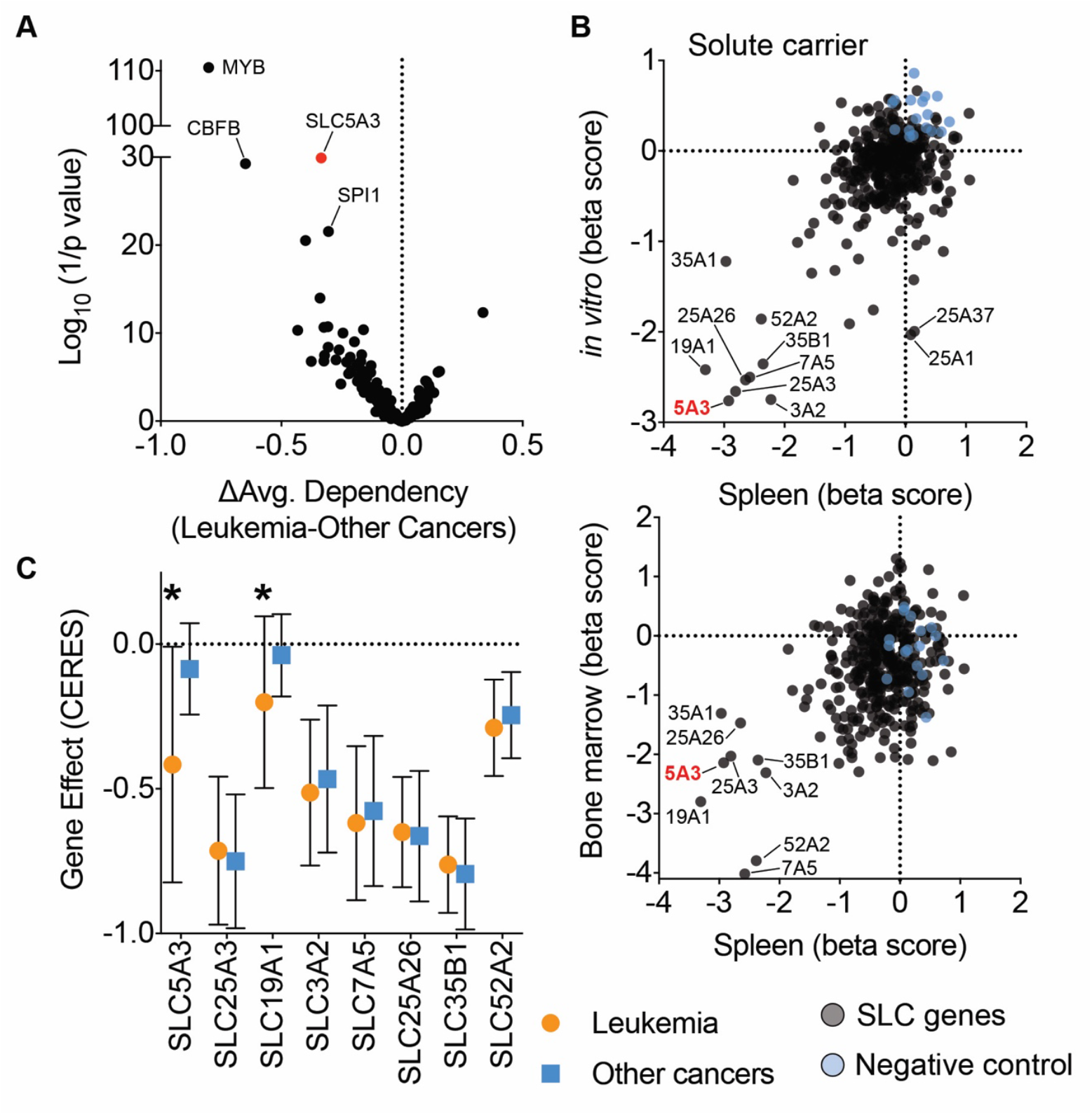
Dependency on SLC5A3 is *in vivo*-relevant and AML-specific. (A) Analysis of Project Achilles genetic screening data, evaluating the leukemia-biased essentiality of the *in vivo*-relevant dependencies. The *in vivo* screening dependencies were determined by *p*-value <0.05, FDR <0.25, beta score <0. The average gene effect scores were calculated using Achilles Gene Effect 20Q3 data. The *p* values compared gene effect scores in leukemia vs. other cancers using a unpaird Student’s t-test. (B) Summary of *in vivo* vs *in vitro* CRISPR screening results of SLC sgRNA library. (C) Comparison of dependencies of top SLC candidates comparing leukemia with other type of cancers. The dependency of SLC5A3 is the most leukemia-biased among the candidates. Data is obtained from DepMap database (20Q3). Plotted as average gene effect score with standard deviation in leukemia cell lines vs. non-leukemia cell lines.

**Figure S4.**
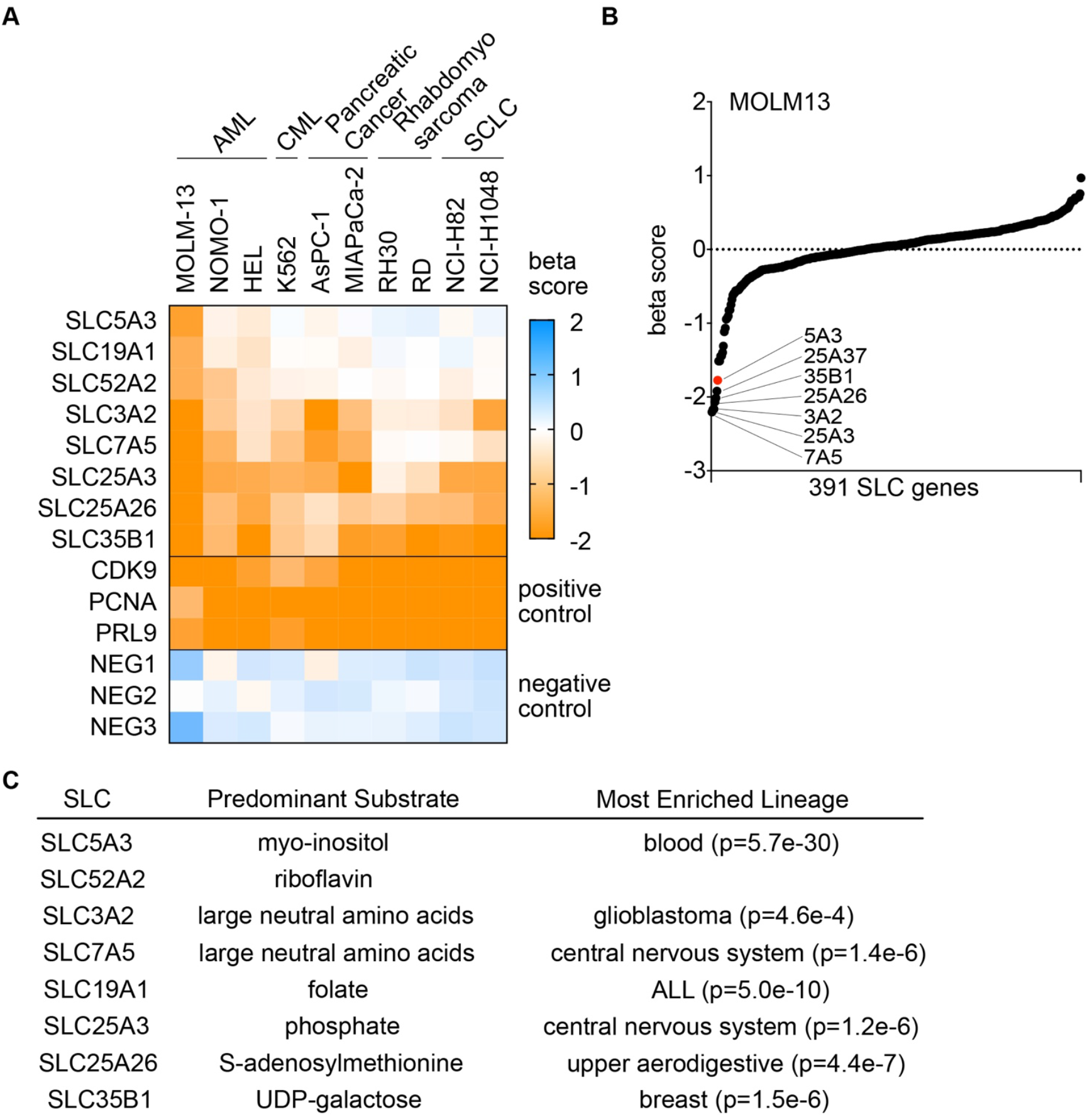
Top SLC targets identified from CRISPR screening in leukemia. (A) Heatmap of beta scores from *in vitro* CRISPR screening in the indicated cancer cell lines. (B) Ranking of top SLC dependencies from *in vitro* screening in MOLM-13 cells. (C) Top SLC candidates and their predominant substrates and enriched lineage.

### Validation of SLC5A3 dependency in a subset of AML cell lines

To validate SLC5A3 as a dependency in AML, we performed competition-based proliferation assays following lentiviral transduction with sgRNA-expressing vectors in Cas9-expressing AML cell lines. The genome editing efficiency of these sgRNAs targeting SLC5A3 was validated using RT-qPCR and Surveyor assays (Figure S5A). Using this approach, we validated that the cell fitness of several AML lines was impaired by SLC5A3 sgRNAs (e.g. MOLM-13, SEM, MV4-11, U937, EOL1), whereas other AML lines were SLC5A3 independent (e.g. HEL, KASUMI, SET-2) (Figure 3A, Figure S5B-C). As a control, an sgRNA targeting the DNA replication protein PCNA suppressed the growth of all cancer cell lines tested. cDNA rescue experiments also supported that these cell fitness alterations were due to a requirement for the *SLC5A3* gene product and rule out off target effects as contributing to this phenotype (Figure S6). We also verified that knockout of *SLC5A3* suppressed the growth of MOLM-13 cells under *in vivo* conditions (Figure 3B-E; Figure S7A-C). Collectively, these findings demonstrate that SLC5A3 is essential in a subset of AML cell lines under both *in vitro* and *in vivo* conditions.

**Figure 3.**
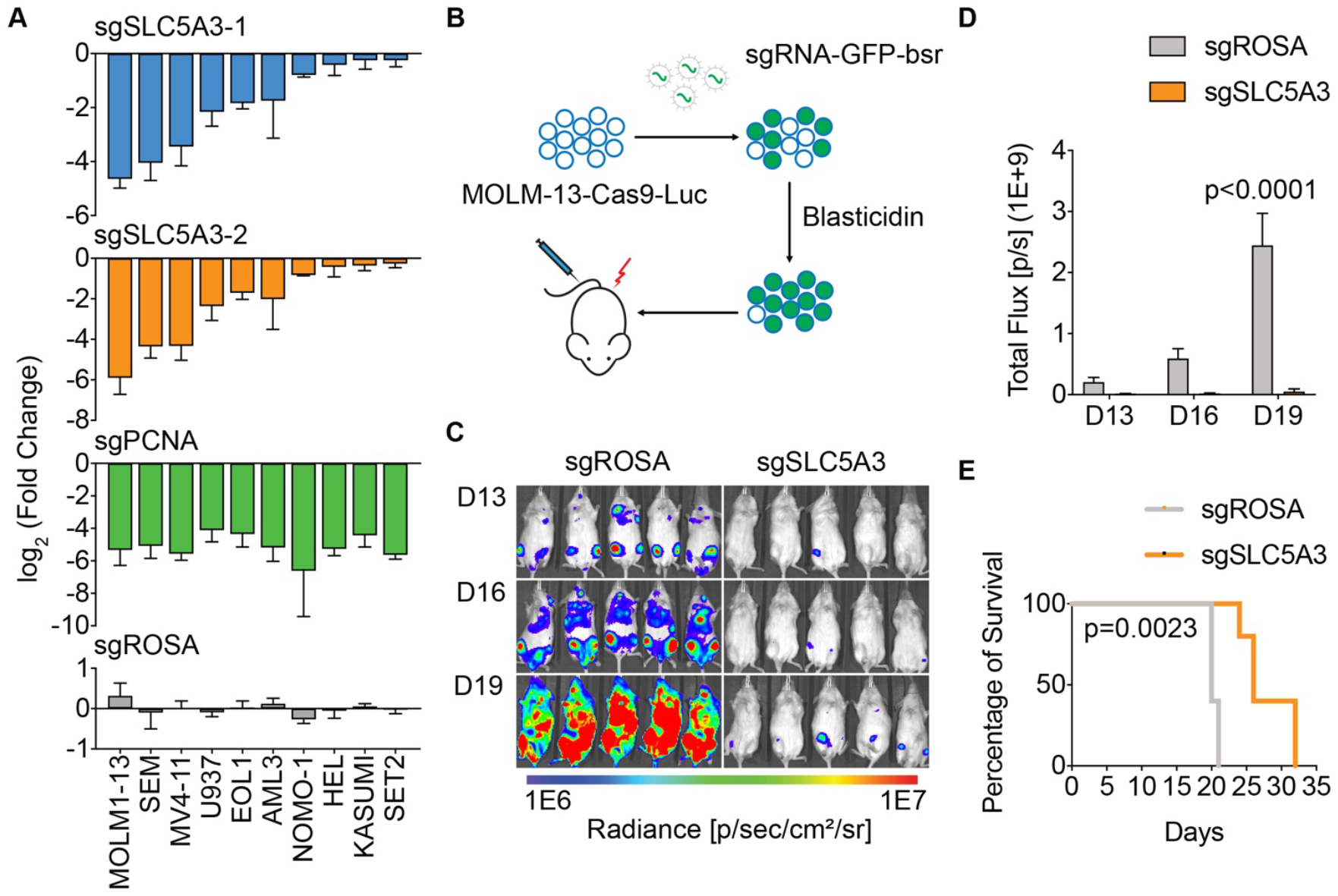
Validation of SLC5A3 dependency in a subset of AML cell lines. (A) Summary of *in vitro* validation experiments of individual sgRNAs (linked to GFP) in competition-based proliferation assays in AML cell lines. Plotted is the log2 transformed fold change of GFP positive cell percentage at day 18 vs. day 3 post the infection of sgRNA lentivirus. Shown as mean with standard deviation of 2~7 biological replicates. (B) Procedure of *in vivo* validation of SLC5A3 dependency in human AML. MOLM-13 cells expressing Cas9 and Luciferase were transduced with sgROSA or sgSLC5A3-2 (linked to GFP/bsr), and then selected with Blasticidin. The SLC5A3 knock-out cells (sgSLC5A3) and control cells (sgROSA) were then transplanted into NSG mice via tail vein injection. (C) Bioluminescence imaging of NSG mice transplanted with MOLM-13 cells transduced with the indicated sgRNA. The images were taken on day 13, 16, and 19 post-transplantation. (D) Quantification of bioluminescence from (C). Values represent total flux as photons per second (Mean ± Standard Deviation). The *p*-value was calculated using unpaired Student’s t test (n=5). (E) Survival curves of mice shown in (C). The *p*-value was calculated using Log-rank (Mantel-Cox) test (n=5).

**Figure S5.**
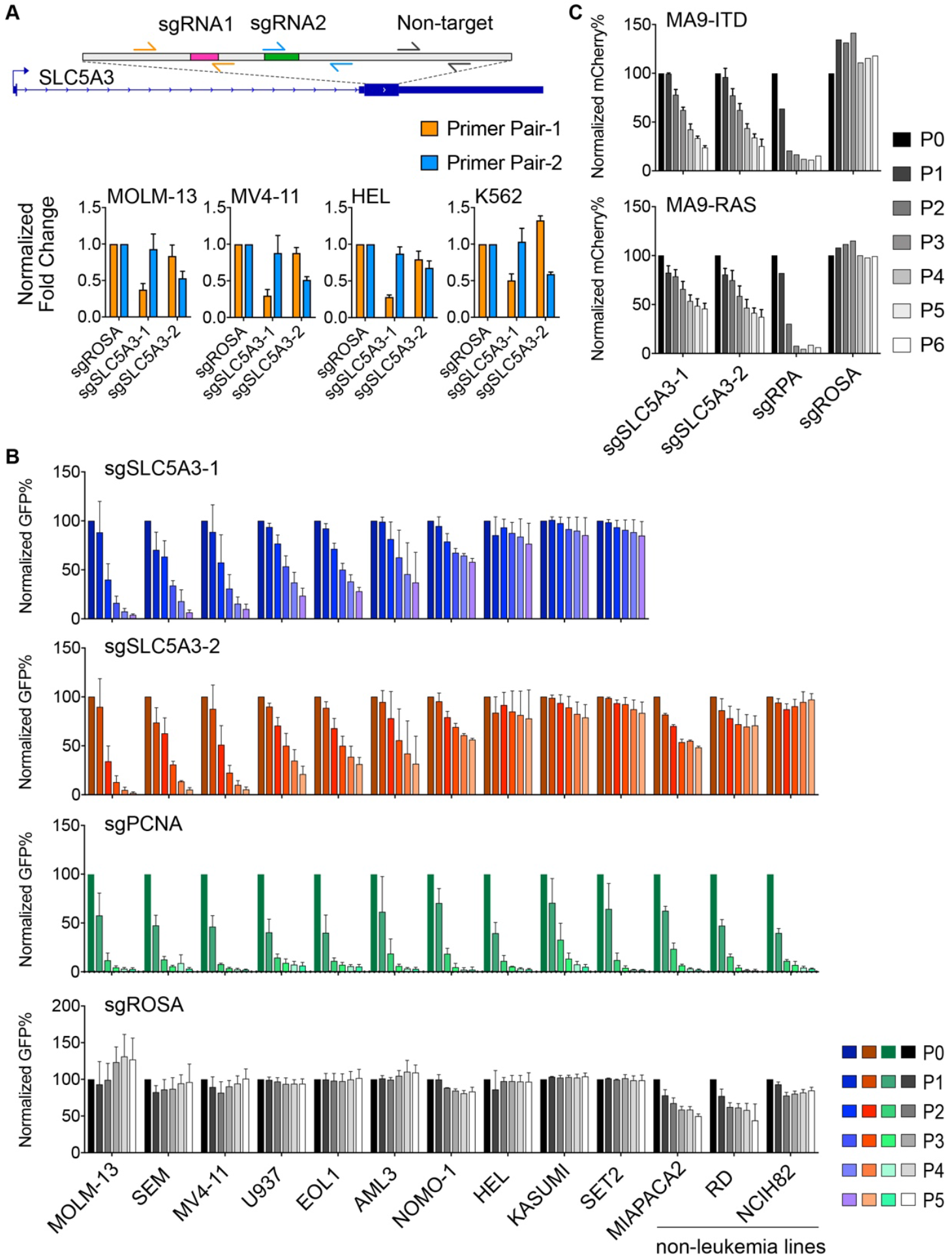
Validation of SLC5A3 dependency in AML cell lines. (A) RT-qPCR shows genomic editing in sgRNA targeted regions. (B) Competition-based proliferation assays in AML cell lines and non-leukemia cell lines infected with indicated sgRNA linked to GFP. Related to Figure 3A. (C) Competition-based proliferation assays in engineered human AML lines MA9-ITD and MA9-RAS infected with indicated sgRNA linked to mCherry.

**Figure S6.**
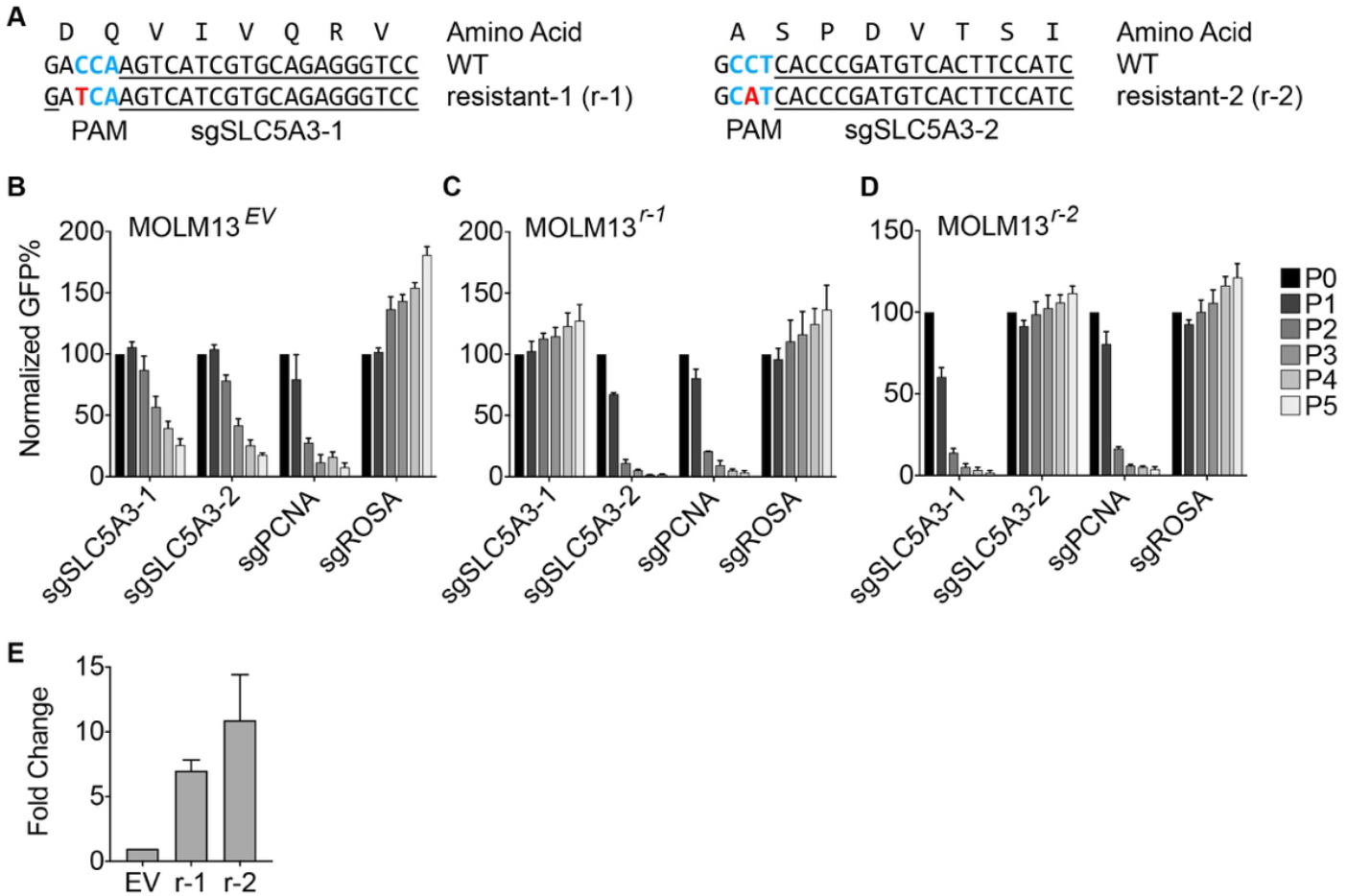
Rescue of SLC5A3 sgRNA with sgRNA-resistant SLC5A3 cDNA. (A) Design of sgRNA-resistant SLC5A3 cDNA. (B-D) Competition-based proliferation assays following transduction of indicated sgRNAs in Cas9 expressing MOLM-13 cells that are infected with EV (B), r-1 (C), or r-2 (D). (Shown as mean with standard deviation of 3 replicates with independent infections). (E) RT-qPCR shows overexpression of sgRNA resistant SLC5A3 cDNA (r-1, r-2).

**Figure S7.**
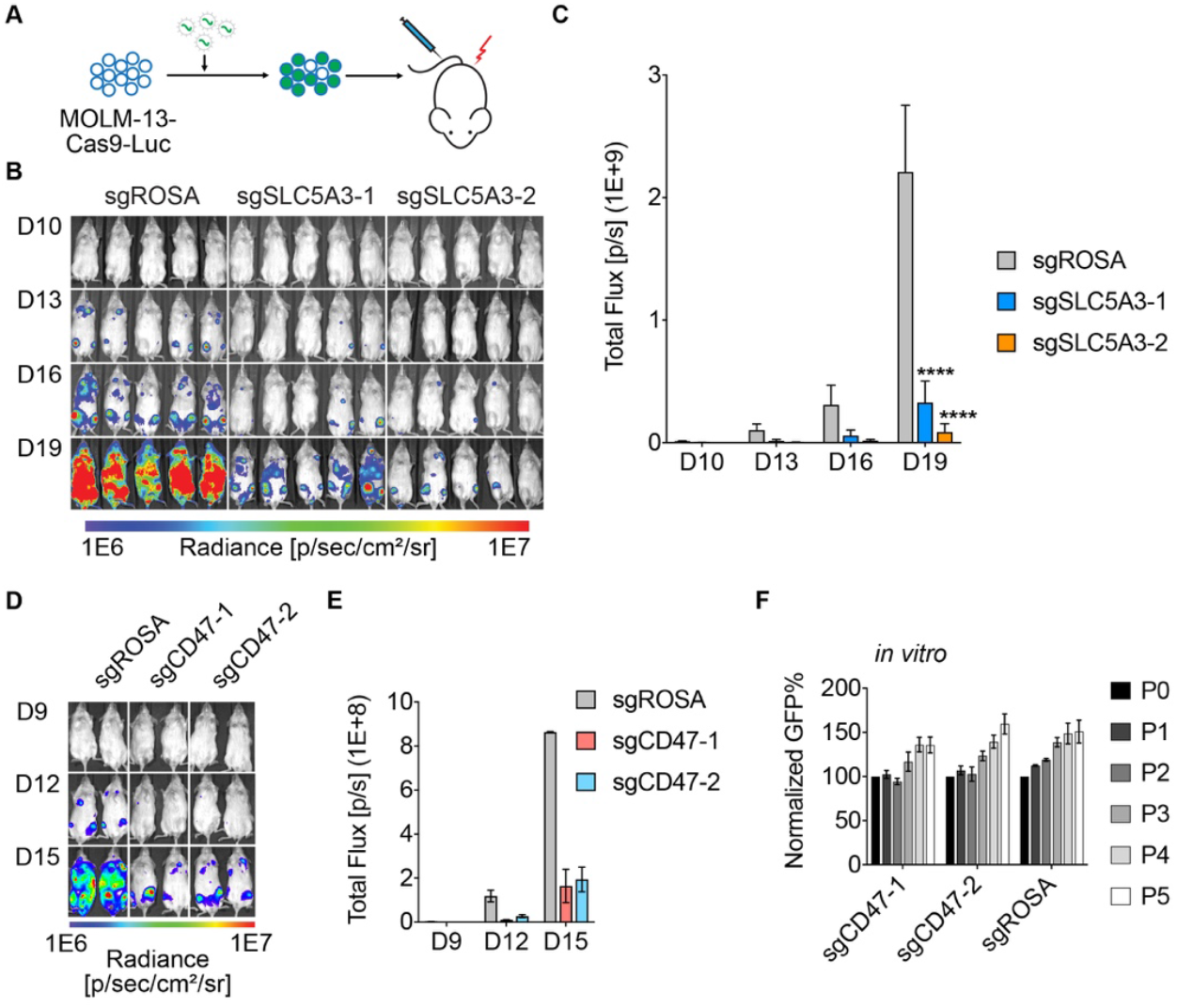
Validation of SLC5A3 and CD47 dependency in AML *in vivo.* (A) Procedure for *in vivo* validation of SLC5A3 or CD47 dependency in human AML. MOLM-13 cells expressing Cas9 and Luciferase were transduced with indicated sgRNA (linked to GFP) with high infection rate (>90%). The cells were then transplanted to NSG mice via tail vein injection. (B) Bioluminescence imaging of NSG mice transplanted with MOLM-13 cells transduced with the indicated sgRNA. The images were taken on day 10, 13, 16, and 19 post-transplantation. (C) Quantification of bioluminescence from (B). Values represent total flux as photons per second (Mean ± Standard Deviation). The *p*-value was calculated using unpaired Student’s t test (n=5). (**** *p*<0.0001). (D) Bioluminescence imaging of NSG mice transplanted with MOLM-13 cells transduced with the indicated sgRNA. The images were taken on day 9, 12, and 15 post-transplantation. (E) Quantification of bioluminescence from (D). Values represent total flux as photons per second. (F) Competition-based proliferation assays in Cas9 expressing MOLM-13 cells transduced with indicated sgRNA linked to GFP.

### SLC5A3-dependent AML lines are myo-inositol auxotrophs

Since the primary substrate for SLC5A3 is myo-inositol (Schneider 2015), we performed liquid chromatography mass spectrometry (LC-MS) to measure intracellular myo-inositol levels in cancer cell lines following SLC5A3 inactivation. As expected, knockout of SLC5A3 in MOLM-13 or MV4-11 cells led to a significant reduction of intracellular myo-inositol, whereas levels of other metabolites (e.g. glutamine) were unaffected (Figure 4A). This finding prompted us to evaluate the impact of depleting myo-inositol from the media of AML cell lines. Remarkably, the SLC5A3-dependent cell lines (MOLM-13, MV4-11, U937) arrested their growth following myo-inositol depletion, whereas the growth of SLC5A3-independent lines (HEL, SET-2, and K562) was unaffected by myo-inositol depletion (Figure 4B-C). Taken together, these experiments suggest that SLC5A3 dependency is associated with a myo-inositol auxotrophy in AML.

**Figure 4.**
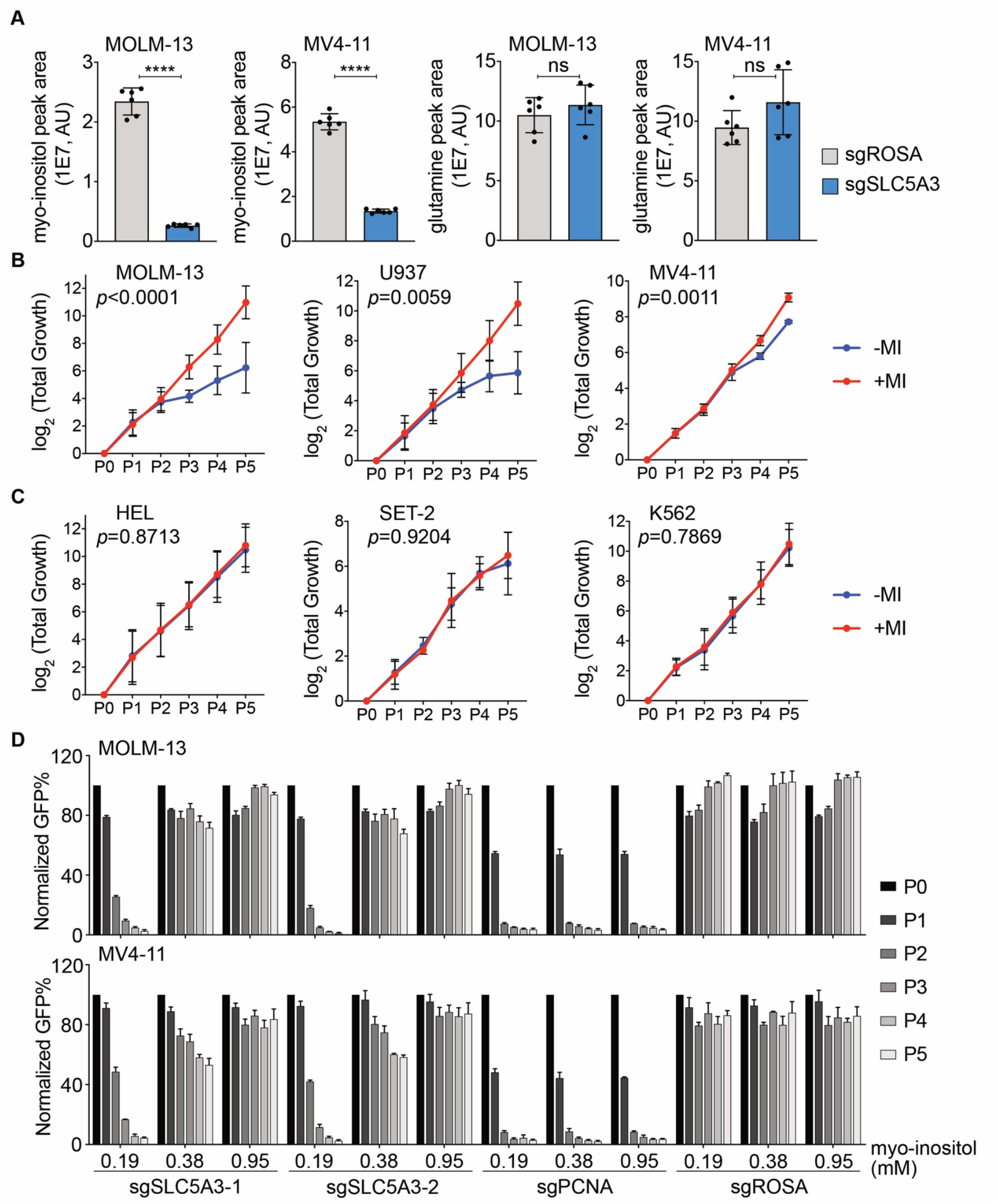
SLC5A3-dependent AML lines are myo-inositol auxotrophs. (A) Liquid Chromatography Mass Spectrometry (LC-MS) analysis of cellular myo-inositol and glutamine levels in MOLM-13 and MV4-11 cells transduced with indicated sgRNA. The *p*-value was calculated using unpaired Student’s t test (n=6). (**** *p*<0.0001). (B-C) Depletion of myo-inositol in the cell culture medium affects the growth of SLC5A3-dependent lines MOLM-13, MV4-11, and U937 (B), but not SLC5A3-independent lines HEL, K562, and SET-2 (C). A customized RPMI cell culture medium without myo-inositol, supplied with dialyzed FBS, was used (-MI). Additional myo-inositol supplement (1.11 mM/0.2 g/L) was added into the medium for rescue (+MI). (Shown as mean with standard deviation of 2~4 biological replicates. The *p* values were calculated using 2way ANOVA Sidak test). (D) Competition-based proliferation assays in MOLM13 and MV4-11 cells transduced with indicated sgRNA, and treated with additional myo-inositol supplement. The base level of myo-inositol in RPMI medium is 0.035 g/L (0.19 mM). To rescue the phenotype, additional myo-inositol was added to the cell culture to make myo-inositol concentration as 0.07 g/L (0.38 mM) and 0.175 g/L (0.95 mM). Myo-inositol from FBS (~0.9 mM, determined by LC-MS, 10% FBS was added to cell culture medium) was not included. (Shown as mean with standard deviation of 3 replicates with independent infections).

It has been found that developmental abnormalities in *Slc5a3^-/-^* mice can be rescued by providing supra-physiological concentrations myo-inositol in the diet, which can be transported into cells in a SLC5A3-independent manner via low affinity transporters (Berry et al. 2003, Chau et al. 2005). This observation motivated us to perform analogous experiments in SLC5A3-dependent AML cell lines by supplementing normal growth media with additional myo-inositol. This revealed that a ~5-fold increase in myo-inositol concentrations in regular growth media (RPMI) was sufficient to bypass the necessity of SLC5A3 in MOLM-13 and MV4-11 cell line contexts (Figure 4D). These findings lend strong support that the essential function of SLC5A3 in AML is the transport of extracellular myo-inositol into the cell interior.

### Transcriptional silencing of ISYNA1 leads to SLC5A3 dependency through a synthetic lethal genetic interaction

We next investigated the mechanism underlying SLC5A3 dependency and myo-inositol auxotrophy in AML. Importantly, SLC5A3 dependency did not correlate with expression levels of SLC5A3, nor did it correlate with silencing of the alternative myo-inositol transporters SLC5A11 and SLC2A13 (Figure S8). Furthermore, we performed triple knockout experiments of SLC5A3, SLC5A11, and SLC2A13, which failed to reveal evidence of redundancy among these three transporters (Figure S9). These results led us to consider whether the myo-inositol auxotrophy in AML was due instead to an acquired defect in *de novo* myo-inositol biosynthesis (Figure 5A). By analyzing the mRNA levels of known myo-inositol biosynthesis genes, we noticed a significant inverse correlation between SLC5A3 dependency and expression of *ISYNA1*, which encodes the rate-limiting enzyme in myo-inositol biosynthesis (Stein & Geige 2002) (Figure 5B). We confirmed the heterogeneous expression of ISYNA1 in AML cell lines using western blotting, which mirrored the heterogeneity of mRNA levels (Figure 5C).

**Figure 5.**
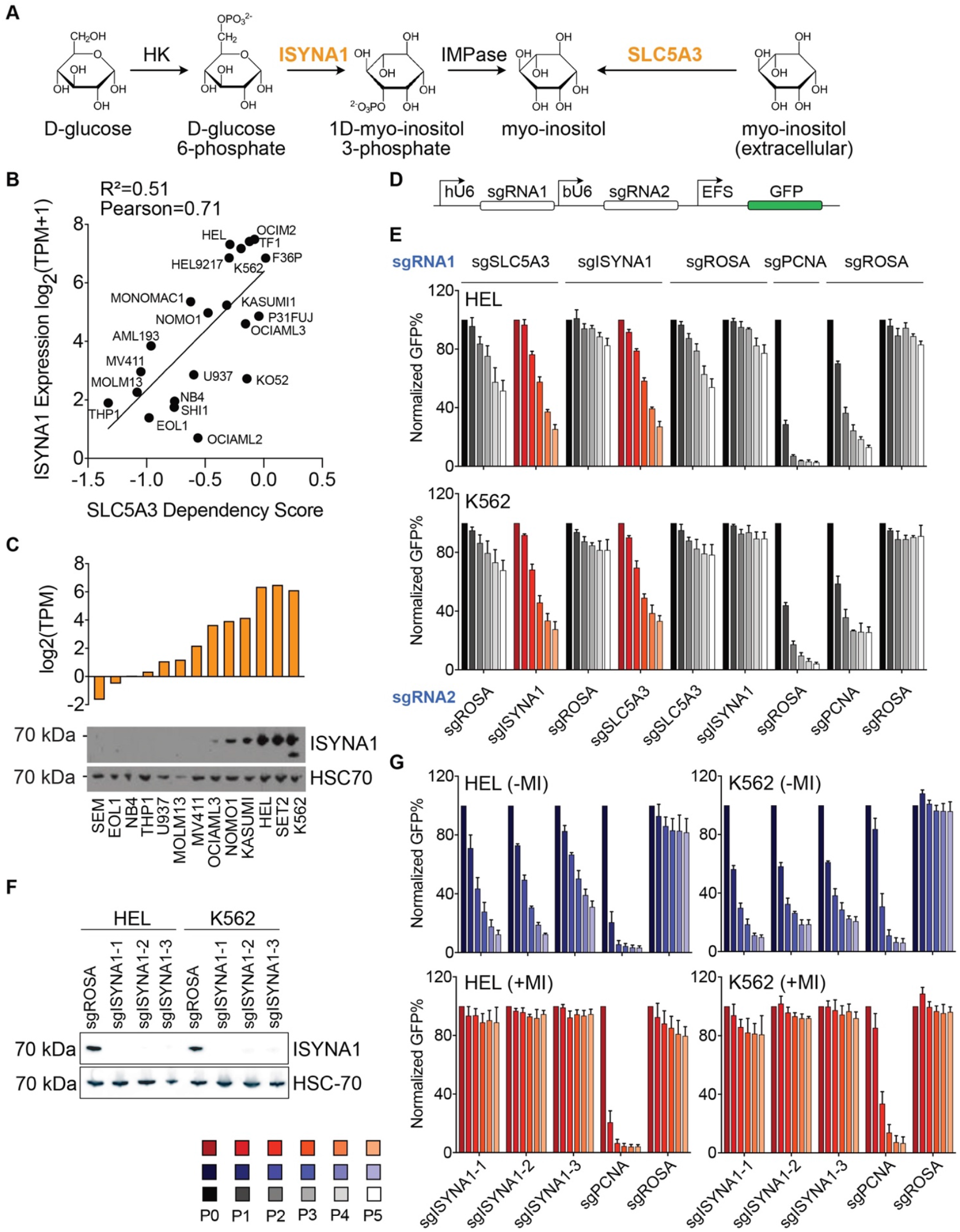
Transcriptional silencing of ISYNA1 leads to SLC5A3 dependency through a synthetic lethal genetic interaction. (A) Summary of two major sources of cellular myo-inositol: transported from extracellular environment by SLC5A3; *de novo* biosynthesis from glucose. The rate limiting de-novo myo-inositol biosynthesis enzyme is inositol-3-phosphate synthase 1, encoded by the *ISYNA1* gene. (B) SLC5A3 dependency in AML (and K562) cell lines correlates with ISYNA1 expression. Data from DepMap database (Achilles 20Q3). (C) Expression of ISYNA1 in AML (and K562) cell lines. The ISYNA1 gene mRNA expression data was obtained from Cancer Cell Line Encyclopedia (CCLE) database. The ISYNA1 protein expression in AML lines and K562 (CML) was detected by Western blot, shown as one of two replicates. (D) A double knock-out construct that co-targets two genes by two sgRNAs driven by human U6 (hU6) or bovine U6 (bU6) promoter, respectively. The sgRNAs are linked to GFP. (E) Competition-based proliferation assays in Cas9 expressing SLC5A3-independent cell lines, following infection with indicated co-targeting sgRNA constructs. Double knock-out both ISYNA1 and SLC5A3 leads to growth phenotype (red). (Shown as mean with standard deviation of 3 replicates of independent infections.). (F) Western blot shows the depletion of ISYNA1 protein in HEL and K562 cells following infection with indicated sgRNA. (G) Competition-based proliferation assays in Cas9 expressing HEL and K562 cells following infection with indicated sgRNA. The experiment was performed in myo-inositol deficient RPMI medium without myoinositol supplement (-MI, blue), or with 0.2 g/L (1.11 mM) myo-inositol supplement (+MI, red). (Shown as mean with standard deviation of 3 replicates with independent infections.)

The findings above led us to hypothesize that a genetic redundancy may exist between ISYNA1 and SLC5A3, which represent two independent strategies for maintaining intracellular myo-inositol levels (Figure 5A). To evaluate this possibility, we targeted *ISYNA1* and *SLC5A3* (alone and in combination) in HEL and K562, which are leukemia lines that express both genes. Competition-based proliferation assays using bi-cistronic sgRNA vectors demonstrated that dual targeting of *ISYNA1* and *SLC5A3* led to a more severe proliferation defect than the loss of each gene individually (Figure 5D-E, Figure S10A). LC-MS analysis demonstrated that a combined deficiency of ISYNA1 and SLC5A3 led a more severe decrease in intracellular myo-inositol than observed in either single gene knockout (Figure S10B). In addition, we found that knockout of *ISYNA1* alone was sufficient to convert HEL and K562 cells into myo-inositol auxotrophs (Figure 5F-G). Taken together, these experiments validate a genetic redundancy between ISYNA1 and SLC5A3 to sustain intracellular myo-inositol and cell viability.

**Figure S8.**
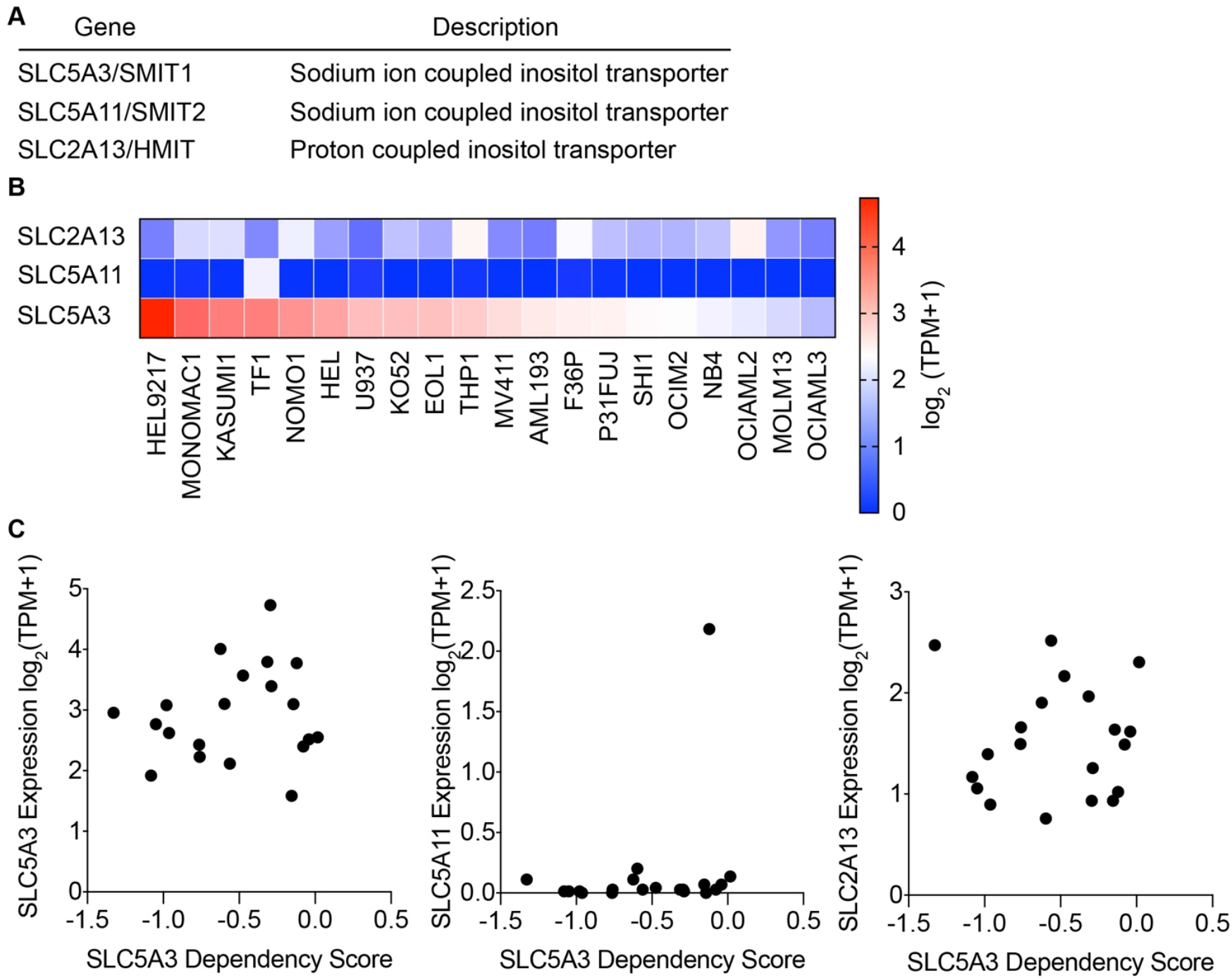
Correlation between SLC5A3 dependency and myo-inositol transporter gene expression. (A) Three myo-inositol transporters in human. (B) Gene expressions of all three myo-inositol transporters in AML cell lines. Data was obtained from CCLE database. (C) Correlation between SLC5A3 dependency and the expression levels of SLC5A3, SLC5A11, or SLC2A13 in AML cell lines. Data was obtained from DepMap database (Achilles 20Q3).

**Figure S9.**
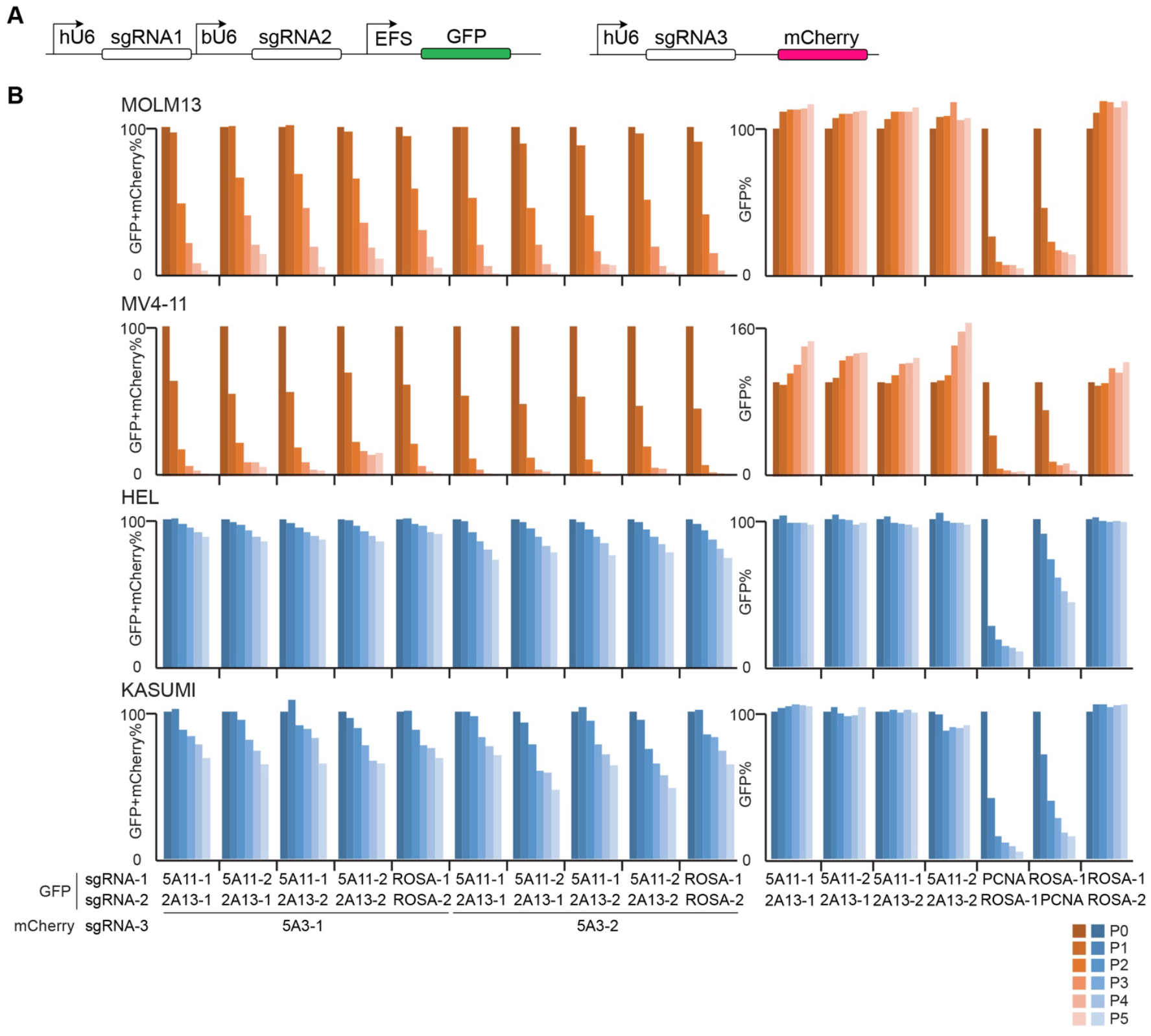
Triple knock-out of all three myo-inositol transporters in AML lines. (A) Overview of triple knock-out by combination of a dual targeting sgRNA construct linked to GFP with a single targeting sgRNA construct linked to mCherry. (B) Competition-based proliferation assays in SLC5A3-dependent AML lines MOLM-13, MV4-11 (orange), and SLC5A3-independent AML lines HEL, KASUMI (blue), following infection with indicated sgRNA constructs.

**Figure S10.**
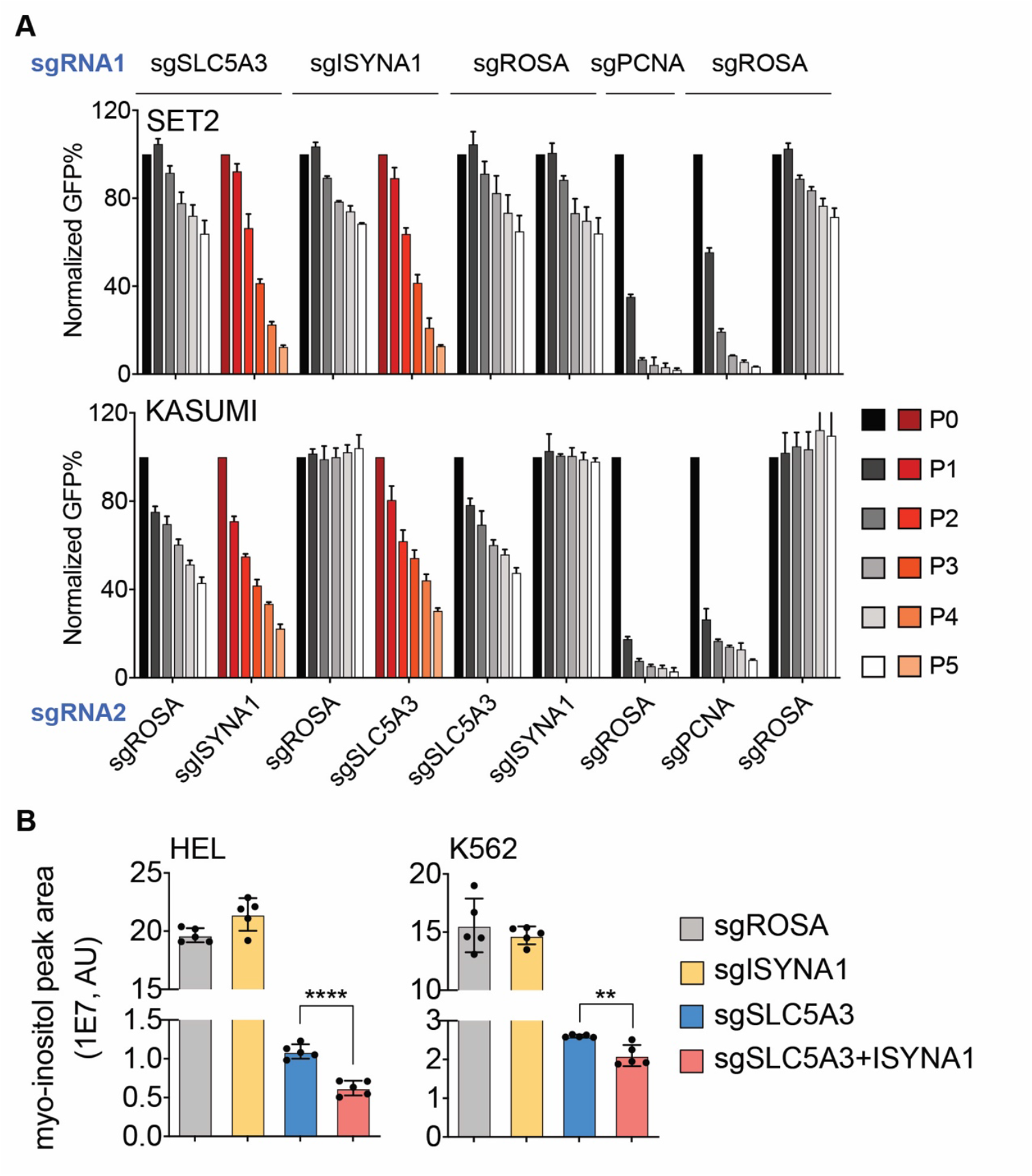
Knock-out of ISYNA1 leads to SLC5A3 dependency and a decrease of cellular myo-inositol. (A) Competition-based proliferation assays in Cas9 expressing SLC5A3-independent cell lines, following infection with indicated co-targeting sgRNA constructs. Double knock-out both ISYNA1 and SLC5A3 leads to growth phenotype (red). (Shown as mean with standard deviation of 3 replicates of independent infections.) (B) LC-MS analysis in Cas9 expressing HEL and K562 cells transduced with indicated sgRNA. The *p*-value was calculated using unpaired Student’s t test (n=5). (**** *p*<0.0001, ** *p*=0.0034)

To validate that ISYNA1 silencing is the underlying cause of myo-inositol auxotrophy and SLC5A3 dependency in AML, we lentivirally expressed a ISYNA1 cDNA in AML cell lines that lacked endogenous ISYNA1 expression (MOLM-13, MV4-11, EOL1, and U937) (Figure 6A). Remarkably, restoring ISYNA1 into each of these contexts was sufficient to bypass the essentiality of SLC5A3 when cultured in normal growth media (Figure 6B). Furthermore, forced ISYNA1 expression also eliminated the myo-inositol auxotrophy by restoring intracellular concentrations of this metabolite (Figure 6C-D). These experiments validate that transcriptional silencing of ISYNA1 is the cause of myo-inositol auxotrophy in AML.

**Figure 6.**
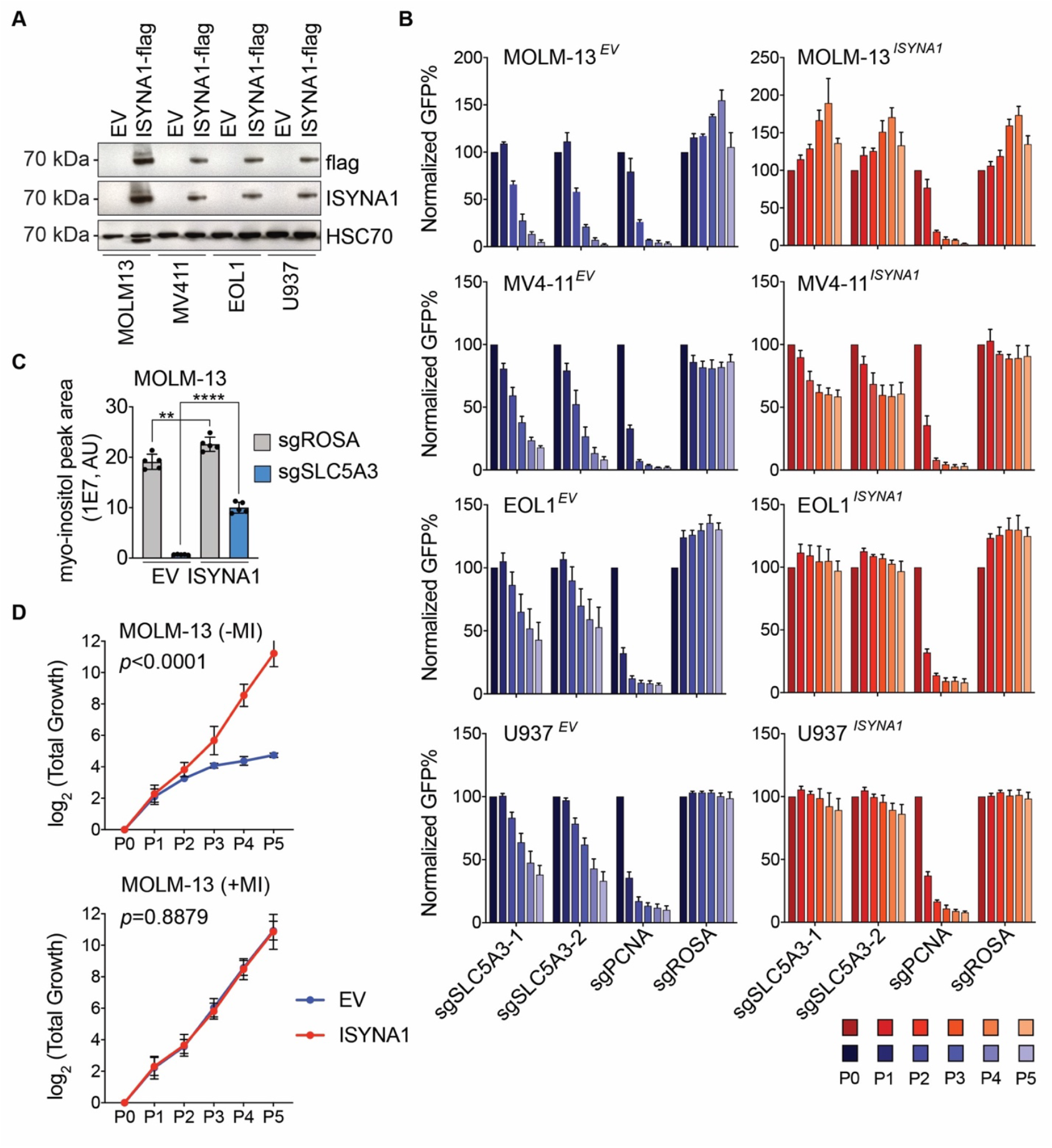
Re-expression of ISYNA1 eliminates SLC5A3 dependency and the myo-inositol auxotrophy. (A) Western blot shows overexpression of flag-tagged ISYNA1 in indicated AML cell lines. (B) Competition-based proliferation assays in SLC5A3-dependent AML lines expressing EV (empty vector, blue) or ISYNA1(red), following infection with indicated sgRNA linked to GFP. (Shown as mean with standard deviation of 3 replicates from independent infections). (C) LC-MS analysis in MOLM-13 cells expressing EV or ISYNA1, following infection with indicated sgRNA. The *p*-value was calculated using unpaired Student’s t test (n=5). (** *p*=0.0054, **** *p*<0.0001). (D) Overexpression of ISYNA1 in MOLM-13 cells rescues the growth defect in myo-inositol deficient medium (-MI), but has no impact on the cell growth in medium with myo-inositol supplement (+MI). (Shown as mean with standard deviation of two biological replicates, each includes 3 replicates with independent infection. The *p* value was calculated using 2way ANOVA Sidak test.)

### Aberrant ISYNA1 DNA methylation associates with IDH mutations in AML patients

We next evaluated whether epigenetic silencing of *ISYNA1* occurs in human AML patient samples. In the cell line AML models, we noticed that transcriptional silencing of *ISYNA1* was associated with DNA hypermethylation and the loss of histone acetylation in the vicinity of its CpG island/promoter (Figure S11A-B). We confirmed DNA hypermethylation at the *ISYNA1* gene in MOLM-13 and MV4-11 lines using Nanopore sequencing (Figure 7A). Notably, these regions remained unmethylated in normal human hematopoietic stem and progenitor cells (HSPC) (Hodges et al. 2011) (Figure 7A, Figure S11C). Having established hypermethylation as a biomarker of ISYNA1 silencing, we next evaluated whether this genomic region was hypermethylated in human AML patient samples analyzed by The Cancer Genomic Atlas (TCGA), in which 170 genetically annotated samples were evaluated using Illumina Infinium 450k DNA methylation assay (Cancer Genome Atlas Research Network et al. 2013). In this study, we found that *ISYNA1* hyper-methylation was present in ~20% of human AML samples, whereas other flanking regions lacked heterogeneity in methylation (Figure 7B, Figure S11D-E). Importantly, the presence of hypermethylation correlated with lower ISYNA1 expression (Figure 7C, Figure S11F). We further evaluated ISYNA1 methylation in an independent cohort of >100 AML patient samples analyzed using reduced representation bisulfite sequencing by Glass et al, which likewise identified hypermethylation within the ISYNA1 CpG island in ~20% of AML cases (Glass et al. 2017) (Figure 7B). In addition, in both AML patient cohorts we found that ISYNA1 hyper-methylation was significantly enriched for patients that possess *IDH1/IDH2* or *CEBPA* mutations (Figure 7B, D). The TCGA AML patient gene expression data further shows patients with IDH2 mutations have significantly lower *ISYNA1* gene expression (Figure 7E). Taken together, these findings validate that aberrant hypermethylaton and silencing of *ISYNA1* occurs recurrently in specific subtypes of AML.

**Figure 7.**
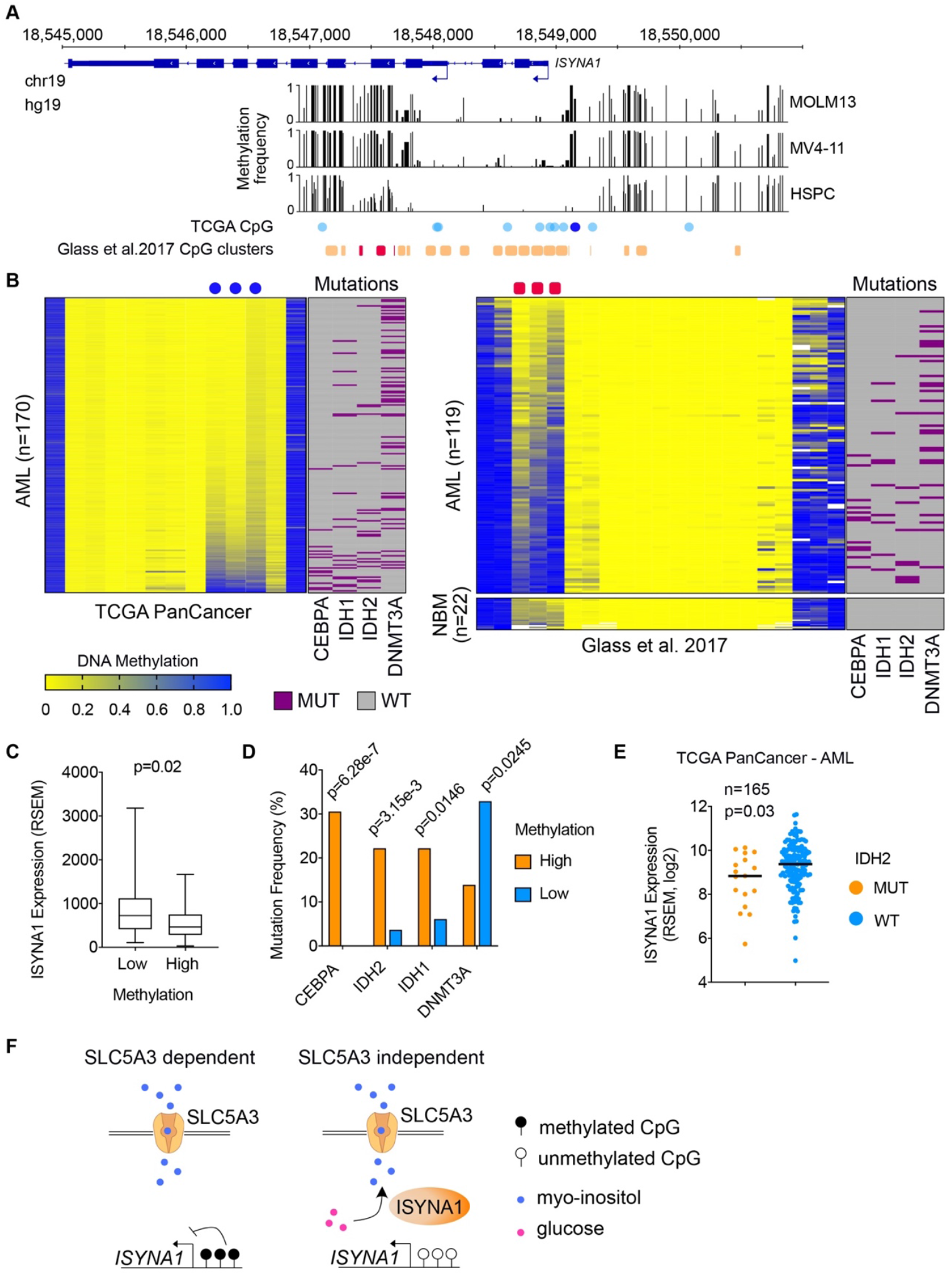
The ISYNA1 gene is epigenetically silenced by aberrant DNA methylation in a subset of AML. (A-B) CpG methylation status at the ISYNA1 gene locus in AML cell lines (MOLM-13, MV4-11), normal hematopoietic stem and progenitor cell (HSPC) (A), and AML patients (B). The AML cell line methylation was measured by Nanopore sequencing. The HSPC methylation was measured by genome-wide bisulfite sequencing, obtained from a previous study (Hodges et al. 2011). The AML patients’ data was obtained from TCGA PanCancer AML database (Cancer Genome Atlas Research Network et al. 2013) or from (Glass et al. 2017). The TCGA AML patients’ data used DNA methylation array with reduced CpG representation, shown as ‘TCGA CpG’. The Glass et al data used reduced representation bisulfite sequencing, and the CpGs were binned every 100 bp, shown as ‘Glass et al. 2017 CpG clusters’. The CpGs at ISYNA1 promoter, as well as downstream of an alternative transcription start site show hypermethylated in SLC5A3-dependent/ISYNA1-low AML lines MOLM-13 and MV4-11, but hypomethylated in normal HSPC cells. These regions were also shown aberrantly methylated in a subset of AML patients, but not in normal bone marrow cells (NBM). (C) ISYNA1 gene expression level is significantly lower in high methylation group, compared to low methylation group (*p*=0.02). The high- and low-groups were determined by the Z scores of average DNA methylation levels of the three promoter CpGs (TCGA) described above (Figure S11E). (D) CEBPA, IDH1, and IDH2 mutations are enriched in the high ISYNA1 methylation group, while DNMT3A mutation is enriched in low methylation group. Data was obtained from TCGA PanCancer AML database via cBioportal. (E) AML patients with IDH2 mutations show lower ISYNA1 gene expression (*p*=0.03). Data was obtained from TCGA PanCancer AML database via cBioportal. (F) A model depicting synthetic lethality between SLC5A3 and ISYNA1 in AML cell lines.

**Figure S11.**
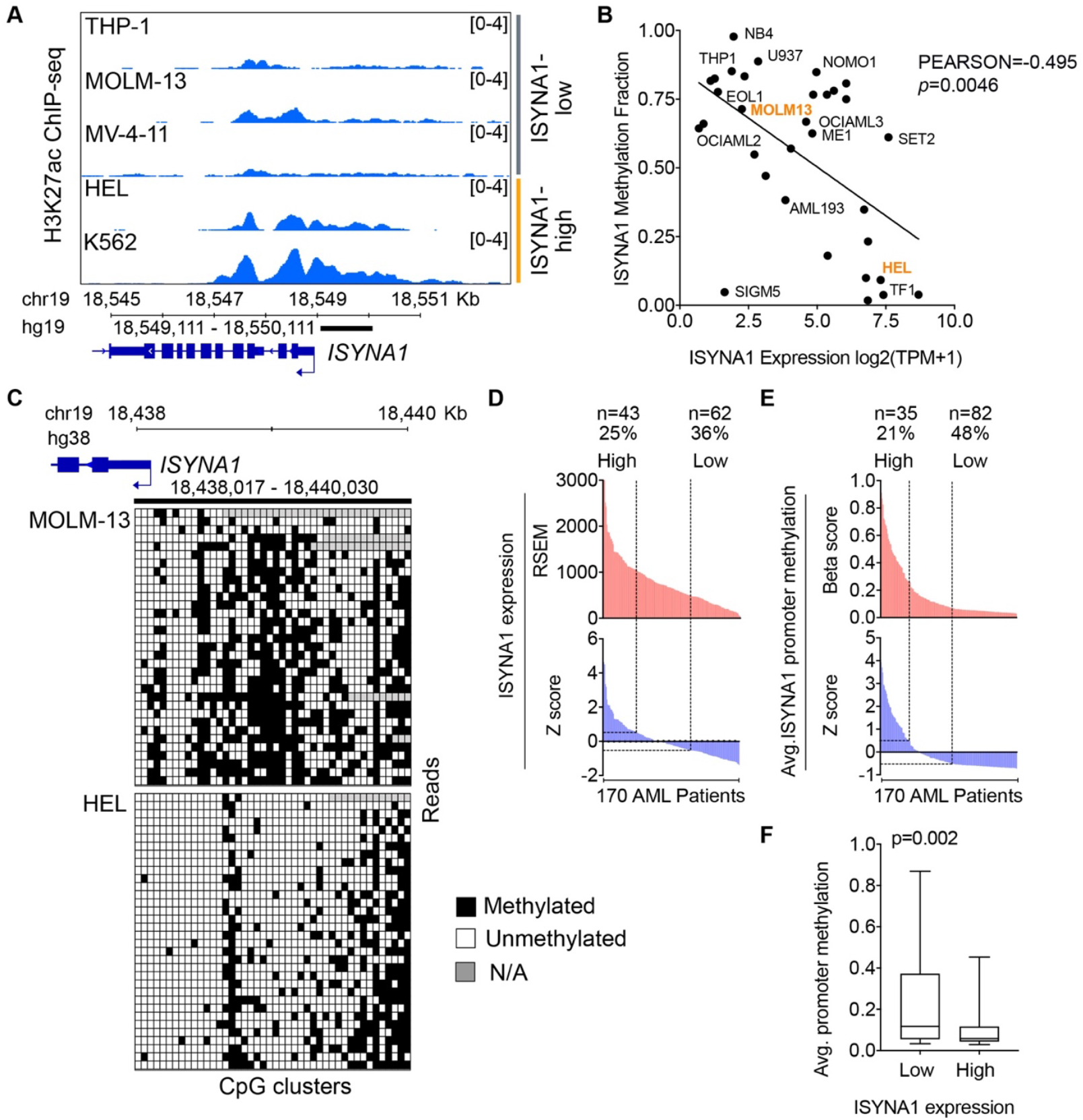
Aberrant DNA methylation at ISYNA1 promoter in AML cell lines and patients. (A) ChIP-seq signal of H3K27ac at the ISYNA1 gene promoter in indicated AML lines and K562. (B) The CpG methylation status in the ISYNA1 gene promoter (1 kb upstream of the transcription start site, chr19 18,549,111-18,550,111, hg19) shows significant correlation with ISYNA1 gene expression in AML cell lines. Data was obtained from DepMap database. (C) Nanopore sequencing at the ISYNA1 gene promoter (chr19 18,438,017-18,440,030, hg38) shows hypermethylated CpGs in SLC5A3-dependent/ISYNA1-low AML line MOLM-13, compared to SLC5A3-independent/ISYNA1-high AML line HEL. The x-axis shows each CpG cluster, and the Y-axis shows each Nanopore sequencing reads from genomic DNA. (D) ISYNA1 gene expression level in AML patients (TCGA PanCancer AML) as shown in both RSEM and Z score. High expression is defined by Z score>0.5, while low expression is defined by Z score<-0.5. (E) ISYNA1 promoter CpG methylation level in AML patients. Shown as the average of the three promoter associated CpGs (Figure 7A-B). High methylation is defined by Z score>0.5, while low methylation is defined by Z score<-0.5. (F) ISYNA1 promoter CpG methylation level is significantly higher in low expression group, compared to high expression group (*p*=0.002).

## Discussion

Our study identifies a myo-inositol auxotrophy as a previously unrecognized metabolic vulnerability in AML. We trace this auxotrophy to aberrant silencing of ISYNA1, which is the rate-limiting enzyme for myo-inositol biosynthesis. Our work suggests that normal cells maintain intra-cellular myo-inositol levels through the partially redundant mechanisms of *de novo* biosynthesis and SLC5A3-dependent transport from the extracellular microenvironment. Our work shows how *ISYNA1* silencing eliminates this redundancy, which in turn drives an elevated demand for SLC5A3-dependent myo-inositol transport in AML (Figure 7F).

The evolution of redundant mechanisms for sustaining intracellular myo-inositol is likely to reflect the vital function of this metabolite, which is well-supported by experiments in diverse eukaryotic species (Livermore et al. 2016). Myo-Inositol is incorporated as head group into a variety of lipids, which function to promote signal transduction in growth-promoting pathways (e.g. AKT-mTOR) (Martelli et al. 2010). Hence, blockade of SLC5A3 in ISYNA1-deficient AML is likely to extinguish AKT-mTOR signaling, which would be expected to cause an acute arrest in cell proliferation. Targeting of SLC5A3 would also be expected to deplete the cell of inositol phosphates, which are known to regulate calcium release as well as the function of chromatin remodeling complexes (Steger et al. 2003, Livermore et al. 2016). These myriad functions of myo-inositol for cell viability are likely to all contribute to the fitness defects of targeting SLC5A3 in AML.

Our study adds to a growing body of evidence highlighting how epigenetic silencing can alter metabolic pathways and lead to nutrient dependencies. Hypermethylation of the argininosuccinate synthetase (ASS1) gene promoter leads to an arginine dependency in many human cancers (Szlosarek et al. 2006, Nicholson et al. 2009, Delage et al. 2012). Similarly, hypermethylation and silencing of asparagine synthetase (ASNS) has been linked to asparagine auxotrophy in lymphoid leukemia (Peng et al. 2001, Ren et al. 2004, Touzart et al. 2019). In addition, promoter hypermethylation of a rate-limiting cholesterol biosynthetic enzyme, squalene mono-oxygenase (SQLE) leads to a cholesterol auxotrophy in anaplastic large cell lymphoma (Garcia-Bermudez et al. 2019). It remains a mystery as to why the CpG islands of metabolic genes are prone to DNA hypermethylation, as there exists no clear evidence that silencing of these genes is under positive selection in human cancer. Since epigenetic and metabolic regulation is tightly interconnected, it is possible that feedback regulation exists between the chromatin regulatory machinery and metabolite levels. As an example, we observe that ISYNA1-silenced AML is correlated with the presence of *IDH1/IDH2* gain-of-function mutations, which produce 2-hydroxyglutarate which in turn inhibits active DNA demethylation (Ward et al. 2010, Figueroa et al. 2010). Thus, it is possible that the unique metabolic state of AML is the predisposing factor for ISYNA1 silencing as a passenger epigenetic event in this disease.

Several mechanisms have been described by which the *ISYNA1* gene is regulated in mammalian cells to control myo-inositol biosynthesis, which may predispose to gene silencing in AML. For example, the inositol hexakisphosphate kinase 1 (IP6K1) has been found in the nucleus where it acts directly to maintain transcription of *ISYNA1* and prevent DNA hypermethylation (Yu et al. 2016). In addition, prior studies have shown that Musashi2, a known oncogene, regulates ISYNA1 expression in the context of pancreatic cancer (Zhou et al. 2020). The tumor suppressor p53 has also been shown to directly regulate the *ISYNA1* promoter (Koguchi et al. 2016). Our own genetic analysis of human samples suggests that mutations of the C/EBPa transcription factor are associated with *ISYNA1* hyper-methylation, which may influence the methylation state of this gene. Collectively, these prior studies suggest that several cancer-relevant pathways converge in regulating *ISYNA1* expression, which might underlie the aberrant silencing of this gene in AML.

Our work raises the possibility that the acquired dependency on SLC5A3 and extra-cellular myo-inositol in AML might have therapeutic significance. In analogy to the use of asparaginase in lymphoblastic leukemias (Egler et al. 2016), one possibility for therapeutic intervention in ISYNA1-deficient AML would be to suppress plasma levels of myo-inositol using an infusion of catabolic enzymes. MIOX (myo-inositol oxygenase) is an enzyme that oxidizes myo-inositol to D-glucuronic acid (Shiue et al. 2014, Zheng et al. 2018). Another enzyme capable of catabolizing myo-inositol is inositol dehydrogenase from *Bacillus subsilis* (Ramaley et al. 1979, Kouzuma et al. 2001). Alternatively, several solute carriers have been targeted using small molecules (Lin et al. 2015). Thus, an attractive strategy would be to suppress SLC5A3 function. While Slc5a3-null mice die soon after birth due to respiratory failure from abnormal development of peripheral nerves, this phenotype can be rescued by prenatal myo-inositol supplementation. Adult Slc5a3-null mice show a decrease of tissue myo-inositol levels and reduced nerve conduction velocity, which can also be alleviated by myo-inositol supplementation (Chau et al. 2005). Taken together, our findings justify an investigation of myo-inositol lowering agents or SLC5A3 blockade as therapeutic interventions for the elimination of ISYNA1-deficient AML cells.

## Material and Method

### Cell Lines and Culture

MOLM-13, MV-4-11, SEM, OCI-AML3, EOL-1, U937, NOMO-1, HEL, SET-2, KASUMI-1 (human acute myeloid leukemia, AML), K562 (human chronic myeloid leukemia, CML), AsPC-1, MIA PaCa-2 (human pancreatic cancer) and NCI-H82 (human small cell lung cancer, SCLC) cells were cultured in RPMI supplemented with 10% fetal bovine serum (FBS). MA9-ITD, MA9-RAS (engineered human AML) cells were cultured in IMDM supplemented with 20% FBS (Mulloy et al. 2008, Wunderlich et al. 2013). RH30, RD (human rhabdomyosarcoma), and HEK293T cells were cultured in DMEM supplemented with 10% FBS. NCI-H1048 (human SCLC) was cultured in DMEM:F12 supplemented with 0.005 mg/mL insulin, 0.01 mg/mL transferrin, 30 nM sodium selenite, 10 nM hydrocortisone, 10 nM β-estradiol, 4.5 mM L-glutamine, and 5% FBS. Penicillin/streptomycin was added to all cell culture. All cell lines were cultured at 37°C with 5% CO2, and were periodically tested negative for mycoplasma contamination.

To make myo-inositol deficient cell culture (-MI), customized RPMI medium without myo-inositol (Bio-Techne) was supplemented with dialyzed FBS (#26400044, ThermoFisher Scientific) and penicillin/streptomycin. The dialyzed FBS shows 100 times less myo-inositol level compared to regular FBS, validated by LC-MS. To measure cell growth in (-MI) cell medium as shown in Figure 4, cells were washed twice with PBS to remove residue cell medium, and then resuspended in (-MI) medium with one million cells per mL. One mL cells were transferred into one well in 12-well plates, with three replicates for each condition. Cell number and viability was measured using automated cell counter (Invitrogen Countess) every two days. For each passage, 200 uL of cells were passed to a new well in 12 well plates containing fresh (-MI) medium. For (+MI) cell medium, 0.2 g/L (1.11 mM) myo-inositol (#I5125, Sigma) was added into the (-MI) medium.

### Animals

*NOD .Cg-Prkdc^scid^Il2r^tm1Wjl^* Tg(CMV-IL3,CSF2,KITLG)1Eav/MloySzJ (known as NSGS) mice expressing human IL3, GM-CSF, and SCF were used for *in vivo* CRISPR screening. <u>NOD.Cg-*Prkdc^scid^Il2rg^tm1Wjl^/SzJ*</u> (known as NSG) mice were used for validating individual gene knock-out *in vivo.* All animals were purchased from The Jackson Laboratory (jax.org). Animal procedures and studies were conducted in accordance with the institutional animal care and use committees at Cold Spring Harbor Laboratory.

### Plasmid Construction

The lentiviral Cas9 vector (LentiV_Cas9_puro/Addgene #108100), the lentiviral sgRNA vectors (LRG2.1-GFP/Addgene #108098, LRG2.1-mCherry/Addgene #108099, the lentiviral luciferase vector (LentiV_Neo_Luc/Addgene #105621), and the lentiviral cDNA expression vector (LentiV_Neo/Addgene #108101) have been described in previous studies (Tarumoto et al. 2018, Tarumoto et al. 2020). DNA oligos of the sgRNAs were cloned into LRG2.1-GFP/mCherry vectors using a BsmBI restriction site. The cDNAs of ISYNA1 (#OHu18552, GenScript) and SLC5A3 (#53836, Addgene) were cloned into LentiV_Neo vector using In-Fusion cloning system (Takara Bio), and flag tag was inserted to the C-terminus of the cDNAs. The same system was used to mutate the PAM sequences in the SLC5A3 gene. Dual targeting vectors were generated by inserting sgRNA1-scaffold-bU6-sgRNA2 sequence (synthesized as gene-blocks by IDT) into the BsmBI site of LRG2.1-GFP-P2A-BlastR vector using Gibson Assembly (NEB). The scaffold sequence for sgRNA1 was generated by replacing the stem-stem loop region of the LRG2.1 scaffold (Shi et al. 2015) with a previously described CRISPRi sgRNA scaffold (Adamson et al. 2016).

### Construction of Domain-Focused sgRNA Library

The Kinase sgRNA library and Transcription factor sgRNA library were described in the previous studies (Tarumoto et al. 2018, Lu et al. 2018). The other sgRNA libraries were generated in a similar fashion. The gene lists were obtained from HUGO Gene Nomenclature Committee database (HGNC, genenames.org). The domain-focused sgRNA libraries were designed by using CSHL CRISPR sgRNA design tool (crispr.cshl.edu), and at least 6 sgRNAs were selected for each gene or domain. Non-targeting sgRNAs as negative control, and sgRNAs targeting essential genes as positive control, were added into the pooled sgRNA library. The pooled library was synthesized on an array platform (Twist Bioscience) and then cloned into the BsmBI restriction site of LRG2.1-GFP vector using Gibson Assembly System (NEB).

### Lentivirus Transduction

Lentivirus was produced in HEK293T cells as described in the previous study (Tarumoto et al. 2018). Briefly, 10 ug of plasmid DNA was mixed with helper plasmids (5 ug VSVG and 7.5 ug psPAX2) and 1 mg/mL polyethylenimine (PEI 25000). The mix was then used to transfect HEK293T cells in 10 cm dish. The mix was replaced by fresh cell media 6 hours post the transfection, and lentivirus-containing supernatant was collected at 24, 48, and 72 hours post-transfection. To infect cells with lentivirus, the target cells were mixed with the virus and 8 ug/mL polybrene in 12 well plates, and then centrifuged at 1700 rpm for 45~60 min. Cell media was changed 24 hours post infection. For cells infected with LentiV_Neo or LentiV_Blast vector, G418 or Blasticidin was added to the cell culture at day 3 post infection, respectively.

### Pooled *in vivo* CRISPR Screening

Cas9 and Luciferase (Luc) expressing MOLM-13 cell line (MOLM-13-Cas9-Luc) was established by infecting MOLM-13 cells with LentiV_Cas9_puro and LentiV_Neo_Luc, and then selected with puromycin and G418. To improve *in vivo* engraftment efficiency, 5 million MOLM-13-Cas9-Luc cells were transplanted into two sub-lethally irradiated (2.5 Gy), 8-week-old female NSG mice through tail vein injection. The cells were collected from the bone marrow of the injected mice ten days post injection by flushing the tibia with phosphate buffered saline (PBS). The collected cells were then cultured in RPMI supplemented with 10% FBS, puromycin and G418 to remove murine cells.

To determine virus titer for the sgRNA library, MOLM-13-Cas9-Luc cells were infected with different volumes of the virus using spin infection technique as described above. Flow cytometer (Guava easyCyte, Millipore) was used to determine the infection rate by measuring the percentage of GFP positive cells. The amount of virus that gave ~30% infection rate was used for further infection experiments.

To ensure at least 1000 fold coverage of each sgRNA in the library, the amount of cells to be infected was calculated using the equation: cell number=sgRNA number/infection rate*1000. For instance, given the sgRNA number in the SLC sgRNA library was ~3000, and the infection rate was ~30%, at least 10 million cells were used for one CRISPR screening. Ten mice were planned to be injected, thus 100 million cells were used for the initial infection.

To perform *in vivo* CRISPR screening, the pre-engrafted MOLM-13-Cas9-Luc cells were infected with the library lentivirus. Three days post infection, the infection rate was checked by flow cytometer to ensure ~30% GFP positive cells. For the SLC sgRNA library, the infected wells were split into three groups. Group 1, ten million cells were spin-collected and kept frozen as P0. Group 2, ten million cells were kept growing in cell culture as *in vitro* CRISPR screening sample, in parallel *of in vivo* CRISPR screening sample. During each passage, the percentage of GFP positive cells was measured, and the cell number was calculated using the equation above to ensure at least 1000 fold coverage of the sgRNA library. Group 3, one hundred million cells were collected, washed with PBS to remove any residual cell media, and then resuspended in 2 mL PBS (5X10^7^ cells/mL). 200 uL of the cells were transplanted to one sub-lethally irradiated (2.5 Gy), 8-week-old female NSGS mouse via tail vein injection. A total number of nine mice were successfully injected. The engraftment and growth of the injected human AML cells was monitored by biofluorescence imaging (IVIS Spectrum System, Caliper Life Sciences). Images were taken 10 minutes after intraperitoneal injection of D-luciferin (50 mg/kg), every 3 days post injection. On day 12 post injection, the animals were euthanized. The spleen (SP) and bone marrow (BM) samples were collected. The spleen was broken down by gently crushing between the frost sides of two glass slides, and then filtered through a cell strainer. The bone marrow was flushed from tibia by injecting with PBS. Meanwhile, the *in vitro* samples were also spin-collected. The genomic DNA from P0, *in vitro* sample, individual SP and BM were purified using QIAamp DNA Mini kit (QIAGEN). The genomic DNA were then diluted into 100 ng/uL. An equal aliquot of genomic DNA from individual SP and BM were pooled to make pooled SP and pooled BM samples. The *in vivo* screening for other sgRNA libraries were performed in a similar fashion, with the amount of cells and the number of animals adjusted according to the sgRNA library size to ensure at least 1000X coverage.

Barcoded sequencing libraries for P0, *in vitro*, 9 individual SP, 9 individual BM, pooled SP, and pooled BM were prepared using the same method as described in the previous study (Tarumoto et al. 2018). The libraries were pooled and analyzed by paired-end sequencing using Miseq (Illumina) with MiSeq Reagent Kit v3 (Illumina).

The sequencing data was de-barcoded and subsequently mapped to the reference sgRNA library using a customized tool as described in the previous study (Shi et al. 2015). The screening data was analyzed using MAGeCK-MLE (Li et al. 2014), and a beta score was calculated for each gene and negative control. Negative beta score indicated negative selection, and positive beta score indicated positive selection. To calculate beta score for the non-targeting negative controls, every 6 negative controls were pooled as a single target for the MAGeCK-MLE calculation. The data was visualized using Prism 8 program.

### Pooled *in vitro* CRISPR Screening

Ten cell lines representing five different type of cancers were used for in vitro CRISPR screening of the SLC sgRNA library, including MOLM-13, HEL, NOMO-1 (AML), K562 (CML), AsPC-1, MIA PaCa-2 (pancreatic cancer), NCI-H82, NCI-H1048 (SCLC), RH30, RD (rhabdomyosarcoma). The Cas9 expressing cell lines were established by infecting with LentiV_Cas9_puro, and then selected with puromycin. The virus titer for each cell line was determined individually as described above. To ensure 1000 fold coverage of the library, at least 10 million cells were initially infected with the library with ~30% infection rate. Day 3 post infection, 10 million cells were collected and kept frozen as P0. Cells were passed every three days, cell number and GFP% were checked every passage. Cells collected at passage 5 and 8 were saved as P5 and P8 samples. Genomic DNA extraction, sequencing library preparation, and data analysis were performed the same way as described above.

### Competition-Based Cell Proliferation Assay

Cas9 expressing cells were infected with sgRNA cloned into LRG2.1-GFP/mCherry, or dual-targeting vector. The percentage of GFP or mCherry positive cells was first measured at day 3 post infection (P0) by flow cytometer (Guava easyCyte, Millipore), and then measured every 2~3 days at each passage, till passage 5 or 6. The relative fluorescent% for each passage was calculated using the equation: relative fluorescent%=[fluorescent%]/[P0 fluorescent%] * 100%. Decrease of the fluorescent% indicated the infected (gene knock-out) cells were out-competed by the uninfected (wide type) cells.

### Western Blot

To detect the ISYNA1 expression levels in human AML cell lines, ten million cells were collected and cell palate was resuspended in 500 uL 1X Laemli sample buffer (Bio-Rad). The cell lysate was then boiled for 10 min, separated by SDS-PAGE, followed by transfer to nitrocellulose membrane and immunoblotting with 1:500 diluted mouse monoclonal anti-ISYAN1(C-9) antibody (#sc-271830, Santa Cruz Biotechnology), and then with 1:10000 diluted polyclonal rabbit anti-mouse HRP antibody (#P0260, Agilent Dako). For control, mouse monoclonal anti-HSC70 (B-6) antibody (#sc-7298, Santa Cruz Biotechnology) was used with 1:10000 dilution.

To detect the recombinant ISYAN1-flag, ten million cells were collected two weeks post infection with cDNA vectors followed by G418 selection. Protein sample preparation and immunoblotting was the same as described above. For flag-tagged proteins, 1:5000 diluted mouse monoclonal anti-flag antibody (Sigma) was used, followed by a secondary antibody of rabbit anti-mouse HRP antibody (#P0260, Agilent Dako) with 1:10000 dilution.

### *In vivo* validation experiments

For the *in vivo* validation as shown in Figure 3, pre-engraft MOLM-13-Cas9-Luc cells were infected with LRG2.1-sgROSA-GFP-BlastR or LRG2.1-sgSLC5A3-2-GFP-BlastR. At day 2 post infection, blasticidin was added to the cell culture to select for infected cells. After 3 days selection, the cells were collected, washed with PBS to remove residual media, and resuspended to 1X10^6^ cells/mL. 100 uL cells were transplanted to one sub-lethally irradiated (2.5 Gy), 8-week-old female NSG mouse via tail vein injection. Five mice were injected for each sgRNA. AML development was monitored by biofluorescence imaging (IVIS Spectrum System, Caliper Life Sciences). Images were taken 10 minutes after intraperitoneal injection of D-luciferin (50 mg/kg), on day 13, 16 and 19 post injection.

For the *in vivo* validation as shown in Figure S7, pre-engraft MOLM-13-Cas9-Luc cells were infected with LRG2.1-sgROSA-GFP, LRG2.1-sgSLC5A3-1-GFP, LRG2.1-sgSLC5A3-2-GFP, LRG2.1-sgCD47-1-GFP, or LRG2.1-sgCD47-2-GFP. Excessive lentivirus was used to ensure >90% infection rate. The cells were then transplanted to NSG mice as described above.

### Metabolomics analysis by liquid chromatography coupled to mass spectrometry (LC-MS)

For the LC-MS analysis shown in Figure 4, 0.5 million cells were infected with LRG2.1-sgROSA-GFP-BlastR or LRG2.1-sgSLC5A3-2-GFP-BlastR in 12 well plates with 6 replicates. For the LC-MS analysis shown in Figure 6, MOLM-13 cells expressing EV or ISYNA1 were infected with LRG2.1-sgROSA-GFP-BlastR or LRG2.1-sgSLC5A3-2-GFP-BlastR in 12 well plates with 5 replicates. The infected cells were selected with blasticidin at day 2 post infection, for 3 days. For the LC-MS analysis shown in Figure S10, HEL and K562 cells were infected with indicated single or dual targeting sgRNA vectors with over 90% infection rate, in 12 well plates with 5 replicates, for 5 days. The cells were then collected for sample preparation.

To prepare cells for LC-MS, cells were quickly washed in PBS before adding 1 mL of ice-cold extraction solution (50% methanol, 30% acetonitrile, 20% H_2_O) per four million cells. The cells were then scrapped and the suspension was snap frozen in liquid nitrogen. Samples were agitated using a Thermomixer (Eppendorf) at 1,400 rpm, 4°C for 15 min followed by incubation at −80°C for 1 h. Samples were then centrifuged at 15,000 rpm, 4°C for 10 min. The supernatants were further centrifuged for another 10 min. The supernatants were stored in autosampler vials at −80°C until analysis.

Intracellular extracts from 5~6 independent cell cultures were analyzed for each condition. Samples were randomized in order to avoid bias due to machine drift and processed blindly. LC-MS analysis was performed using a Vanquish Horizon UHPLC system couple to a Q Exactive HF mass spectrometer (both Thermo Fisher Scientific). Sample extracts (5 μL) were injected onto a Sequant ZIC-pHILC column (150 mm × 2.1 mm, 5 μm) and guard column (20 mm × 2.1 mm, 5 μm) from Merck Millipore kept at 45°C. The mobile phase was composed of 20 mM ammonium carbonate with 0.1% ammonium hydroxide in water (solvent A), and acetonitrile (solvent B). The flow rate was set at 200 μl/min with the previously described gradient (Mackay et al. 2015). The mass spectrometer was operated in full MS and polarity switching mode. The acquired spectra were analyzed using XCalibur Qual Browser and XCalibur Quan Browser software (Thermo Fisher Scientific) by referencing to an internal library of compounds.

### Nanopore sequencing

crRNA guides specific to the regions of interest (ROI) were designed as per recommended guidelines described in the Nanopore infosheet on Targeted, amplification-free DNA sequencing using CRISPR/Cas (Version: ECI_S1014_v1_revA_11Dec2018). The crRNA sequences are provided in Supplemental File 2. Guides were reconstituted to 100 μM using TE (pH 7.5) and pooled into an equimolar mix. For each distinct sample, four identical reactions were prepared parallelly using 5μg gDNA each. Ribonucleoprotein complex (RNPs) assembly, genomic DNA dephosphorylation, and Cas9 cleavage were performed as described in Gilpatrick et al. 2019. Affinity-based Cas9-Mediated Enrichment (ACME) using Invitrogen™ His-Tag Dynabeads™ was performed to pulldown Cas9 bound non-target DNA, increasing the proportion of on-target reads in the sample (Iyer et al. 2020). The resultant product was cleaned up using 1X Ampure XP beads (Beckman Coulter, Cat #A63881), eluted in nuclease-free water, and pooled together. The ACME enriched sample was quantified using Qubit fluorometer (ThermoFisher Scientific) and carried forward to the adapter ligation step as described by Iyer et al. Sequencing adaptors from the Oxford Nanopore Ligation Sequencing Kit (ONT, SQK-LSK109) were ligated to the target fragments using T4 DNA ligase (NEBNext Quick Ligation Module E6056). The sample was cleaned up using 0.3X Ampure XP beads (Beckman Coulter, Cat #A63881), washed with long-fragment buffer (LFB; ONT, SQK-LSK109), and eluted in 15 μl of elution buffer (EB; ONT, LSK109) for 30 min at room temperature. Resultant library was prepared for loading as described in the Cas-mediated PCR-free enrichment protocol from ONT (Version: ENR_9084_v109_revH_04Dec2018) by adding 25 μL sequencing buffer (SQB; ONT, LSK109) and 13 μl loading beads (LB; ONT, LSK109) to 12 μl of the eluate. Each sample was run on a FLO-MIN106 R9.4.1 flow cell using the GridION sequencer.

Real time basecalling was performed with Guppy v3.2, and files were synced to our Isilon 400NL storage server for further processing on the shared CSHL HPCC. Nanopolish v0.13.2 (Simpson et al. 2017) was used to call methylation per the recommended workflow. Briefly, indexing was performed to match the ONT fastq read IDs with the raw signal level fast5 data. The ONT reads were then aligned to the human reference genome (UCSC hg38) using minimap2 v2.17 (Li 2018) and the resulting alignments were then sorted with samtools v0.1.19 (Li et al. 2009). Nanopolish call-methylation was then used to detect methylated bases within the targeted regions – specifically 5-methylcytosine in a CpG context. The initial output file contained the position of the CpG dinucleotide in the reference genome and the methylation call in each read. A positive value for log_lik_ratio was used to indicate support for methylation, using a cutoff value of 2.0. The helper script calculate_methylation.py was then used to calculate the frequency of methylation calls by genomic position.

### RT-qPCR

To analyze sgRNA editing of genomic DNA as shown in Figure S5A, cells were infected with LRG2.1-sgROSA-GFP-BlastR, LRG2.1-sgSLC5A3-1-GFP-BlastR, or LRG2.1-sgSLC5A3-2-GFP-BlastR. At day 2 post infection, blasticidin was added to the cell culture to select for infected cells. After 3 days selection, the cells were collected, and genomic DNA was extracted using QIAamp DNA Mini kit (QIAGEN). For qPCR, 20~25 ng of genomic DNA was used in each reaction with indicated primer set. The Ct value of non-sgRNA-target primer was used to normalize the data. To analyze SLC5A3 (r-1 and r-2) expression as shown in Figure S6E, MOLM-13 cells were infected with the indicated cDNA constructs, and were then selected with G418 for two weeks. Total RNA was extracted using TRIzol (ThermoFisher Scientific), and cDNA was synthesized using qScript cDNA SuperMix (Quanta). All qPCR experiments were performed using Power SYBR Green PCR Master Mix (ThermoFisher Scientific) on a QuantStudio Flex Real-time PCR machine (ThermoFisher Scientific).

### AML patient data

TCGA PanCancer AML patient data, including gene expression and mutation, were downloaded via cBioPortal (cbioportal.org) (Cerami et al. 2012, Gao et al. 2013, Hoadley et al. 2018). The same patients’ DNA methylation data was downloaded via NCI Genomic Data Commons (gdc.cancer.gov) (Cancer Genome Atlas Research Network et al. 2013). The Glass et al data from AML (n=119) and normal (n=22) individuals were downloaded from GEO (accession number GSE98350) (Glass et al., 2017). After filtering and normalizing by coverage, a methylBase object containing the methylation information and locations of cytosines that were present in at least 5 samples per condition (meth.min=5) was generated using MethylKit (version 1.9.3) and R statistical software (version 3.5.1). Percent methylation for each CG for each donor, was calculated using the MethylKit ‘percMethylation’ function. Bedtools intersect function was used to determine overlap with CpGi from hg19.

## ACKNOWLEDGMENTS

We thank James C. Mulloy for sharing genetically engineered human AML cell lines. This work was supported by Cold Spring Harbor Laboratory NCI Cancer Center Support grant 5P30CA045508. Additional funding was provided to C.R.V. by the Pershing Square Sohn Cancer Research Alliance, National Institutes of Health grants R01 CA174793 and P01 CA013106, a Leukemia & Lymphoma Society Scholar Award.

## AUTHORSHIP CONTRIBUTIONS

Y.W., S.V.I., A.S.H.C., Z.Y., M.K., E.R.A., O.K., O.E.D., S.P., K.C., S.G., performed experiments and/or analyzed the data; M.E.F., W.R.M, E.H., and C.R.V. supervised the experiments and analysis; Y.W. and C.R.V. wrote the manuscript.

## References

1. Adamson B, Norman TM, Jost M, Cho MY, Nuñez JK, Chen Y, et al. A Multiplexed Single-Cell CRISPR Screening Platform Enables Systematic Dissection of the Unfolded Protein Response. Cell. 2016;167:1867–1882.e21.

2. Bacci M, Lorito N, Ippolito L, Ramazzotti M, Luti S, Romagnoli S, et al. Reprogramming of amino acid transporters to support aspartate and glutamate dependency sustains endocrine resistance in breast cancer. Cell Rep. 2019;28:104–118.e8.

3. Bai X, Moraes TF, Reithmeier RAF. Structural biology of solute carrier (SLC) membrane transport proteins. Mol Membr Biol. 2017;34:1–32.

4. Bansal VS, Majerus PW. Phosphatidylinositol-derived precursors and signals. Annu Rev Cell Biol. 1990;6:41–67.

5. Berridge MJ, Irvine RF. Inositol phosphates and cell signalling. Nature. 1989;341:197–205.

6. Berry GT, Mallee JJ, Kwon HM, Rim JS, Mulla WR, Muenke M, et al. The human osmoregulatory Na+/myo-inositol cotransporter gene (SLC5A3): molecular cloning and localization to chromosome 21. Genomics. 1995;25:507–13.

7. Berry GT, Wu S, Buccafusca R, Ren J, Gonzales LW, Ballard PL, et al. Loss of murine Na+/myo-inositol cotransporter leads to brain myo-inositol depletion and central apnea. J Biol Chem. 2003;278:18297–302.

8. Blazar BR, Lindberg FP, Ingulli E, Panoskaltsis-Mortari A, Oldenborg PA, Iizuka K, et al. CD47 (integrin-associated protein) engagement of dendritic cell and macrophage counterreceptors is required to prevent the clearance of donor lymphohematopoietic cells. J Exp Med. 2001;194:541–9.

9. Brien GL, Remillard D, Shi J, Hemming ML, Chabon J, Wynne K, et al. Targeted degradation of BRD9 reverses oncogenic gene expression in synovial sarcoma. elife. 2018;7.

10. Burger JA, Kipps TJ. CXCR4: a key receptor in the crosstalk between tumor cells and their microenvironment. Blood. 2006;107:1761–7.

11. Cancer Genome Atlas Research Network, Ley TJ, Miller C, Ding L, Raphael BJ, Mungall AJ, et al. Genomic and epigenomic landscapes of adult de novo acute myeloid leukemia. N Engl J Med. 2013;368:2059–74.

12. Cerami E, Gao J, Dogrusoz U, Gross BE, Sumer SO, Aksoy BA, et al. The cBio cancer genomics portal: an open platform for exploring multidimensional cancer genomics data. Cancer Discov. 2012;2:401–4.

13. Chau JFL, Lee MK, Law JWS, Chung SK, Chung SSM. Sodium/myo-inositol cotransporter-1 is essential for the development and function of the peripheral nerves. FASEB J. 2005;19:1887–9.

14. Coady MJ, Wallendorff B, Gagnon DG, Lapointe J-Y. Identification of a novel Na+/myo-inositol cotransporter. J Biol Chem. 2002;277:35219–24.

15. Cramer SL, Saha A, Liu J, Tadi S, Tiziani S, Yan W, et al. Systemic depletion of L-cyst(e)ine with cyst(e)inase increases reactive oxygen species and suppresses tumor growth. Nat Med. 2017;23:120–7.

16. Croze ML, Soulage CO. Potential role and therapeutic interests of myo-inositol in metabolic diseases. Biochimie. 2013;95:1811–27.

17. DeBerardinis RJ, Chandel NS. Fundamentals of cancer metabolism. Sci Adv. 2016;2:e1600200.

18. Delage B, Luong P, Maharaj L, O’Riain C, Syed N, Crook T, et al. Promoter methylation of argininosuccinate synthetase-1 sensitises lymphomas to arginine deiminase treatment, autophagy and caspase-dependent apoptosis. Cell Death Dis. 2012;3:e342.

19. Dempster JM, Rossen J, Kazachkova M, Pan J, Kugener G, Root DE, et al. Extracting Biological Insights from the Project Achilles Genome-Scale CRISPR Screens in Cancer Cell Lines. BioRxiv. 2019;

20. Egler RA, Ahuja SP, Matloub Y. L-asparaginase in the treatment of patients with acute lymphoblastic leukemia. J Pharmacol Pharmacother. 2016;7:62–71.

21. Eisenberg F, Bolden AH. Biosynthesis of inositol in rat testis homogenate. Biochemical and Biophysical Research Communications. 1963;12:72–7.

22. El-Gebali S, Bentz S, Hediger MA, Anderle P. Solute carriers (SLCs) in cancer. Mol Aspects Med. 2013;34:719–34.

23. Figueroa ME, Abdel-Wahab O, Lu C, Ward PS, Patel J, Shih A, et al. Leukemic IDH1 and IDH2 mutations result in a hypermethylation phenotype, disrupt TET2 function, and impair hematopoietic differentiation. Cancer Cell. 2010;18:553–67.

24. Ganapathy V, Thangaraju M, Prasad PD. Nutrient transporters in cancer: relevance to Warburg hypothesis and beyond. Pharmacol Ther. 2009;121:29–40.

25. Gao J, Aksoy BA, Dogrusoz U, Dresdner G, Gross B, Sumer SO, et al. Integrative analysis of complex cancer genomics and clinical profiles using the cBioPortal. Sci Signal. 2013;6:pl1.

26. Garcia-Bermudez J, Baudrier L, Bayraktar EC, Shen Y, La K, Guarecuco R, et al. Squalene accumulation in cholesterol auxotrophic lymphomas prevents oxidative cell death. Nature. 2019;567:118–22.

27. Garcia-Bermudez J, Williams RT, Guarecuco R, Birsoy K. Targeting extracellular nutrient dependencies of cancer cells. Mol Metab. 2020;33:67–82.

28. Gilpatrick T, Lee I, Graham JE, Raimondeau E, Bowen R, Heron A, et al. Targeted Nanopore Sequencing with Cas9 for studies of methylation, structural variants and mutations. BioRxiv. 2019;

29. Glass JL, Hassane D, Wouters BJ, Kunimoto H, Avellino R, Garrett-Bakelman FE, et al. Epigenetic identity in AML depends on disruption of nonpromoter regulatory elements and is affected by antagonistic effects of mutations in epigenetic modifiers. Cancer Discov. 2017;7:868–83.

30. Grevet JD, Lan X, Hamagami N, Edwards CR, Sankaranarayanan L, Ji X, et al. Domain-focused CRISPR screen identifies HRI as a fetal hemoglobin regulator in human erythroid cells. Science. 2018;361:285–90.

31. Hager K, Hazama A, Kwon HM, Loo DD, Handler JS, Wright EM. Kinetics and specificity of the renal Na+/myo-inositol cotransporter expressed in Xenopus oocytes. J Membr Biol. 1995;143:103–13.

32. Hart T, Chandrashekhar M, Aregger M, Steinhart Z, Brown KR, MacLeod G, et al. High-Resolution CRISPR Screens Reveal Fitness Genes and Genotype-Specific Cancer Liabilities. Cell. 2015;163:1515–26.

33. Hauser G, Finelli VN. The biosynthesis of free and phosphatide myo-inositol from glucose by mammalian tissue slices. J Biol Chem. 1963;238:3224–8.

34. Hitomi K, Tsukagoshi N. cDNA sequence for rkST1, a novel member of the sodium ion-dependent glucose cotransporter family. Biochimica et Biophysica Acta (BBA)-Biomembranes. 1994;1190:469–72.

35. Hoadley KA, Yau C, Hinoue T, Wolf DM, Lazar AJ, Drill E, et al. Cell-of-Origin Patterns Dominate the Molecular Classification of 10,000 Tumors from 33 Types of Cancer. Cell. 2018;173:291–304.e6.

36. Hodges E, Molaro A, Dos Santos CO, Thekkat P, Song Q, Uren PJ, et al. Directional DNA methylation changes and complex intermediate states accompany lineage specificity in the adult hematopoietic compartment. Mol Cell. 2011;44:17–28.

37. Holub BJ. Metabolism and function of myo-inositol and inositol phospholipids. Annu Rev Nutr. 1986;6:563–97.

38. Hu K, Li K, Lv J, Feng J, Chen J, Wu H, et al. Suppression of the SLC7A11/glutathione axis causes synthetic lethality in KRAS-mutant lung adenocarcinoma. J Clin Invest. 2020; 130:17526–6.

39. Irvine RF, Schell MJ. Back in the water: the return of the inositol phosphates. Nat Rev Mol Cell Biol. 2001;2:327–38.

40. Iyer SV, Goodwin S, Kramer M, McCombie WR. Abstract 1360: Understanding genetic variation in cancer using targeted nanopore long read sequencing. Molecular and Cellular Biology / Genetics. American Association for Cancer Research; 2020. page 1360–1360.

41. Jaffe N, Traggis D, Das L, Moloney WC, Hann HW, Kim BS, et al. L-asparaginase in the treatment of neoplastic diseases in children. Cancer Res. 1971;31:942–9.

42. Kitamura H, Yamauchi A, Nakanishi T, Takamitsu Y, Sugiura T, Akagi A, et al. Effects of inhibition of myo-inositol transport on MDCK cells under hypertonic environment. Am J Physiol. 1997;272:F267–72.

43. Koguchi T, Tanikawa C, Mori J, Kojima Y, Matsuda K. Regulation of myo-inositol biosynthesis by p53-ISYNA1 pathway. Int J Oncol. 2016;48:2415–24.

44. Kouzuma T, Takahashi M, Endoh T, Kaneko R, Ura N, Shimamoto K, et al. An enzymatic cycling method for the measurement of myo-inositol in biological samples. Clin Chim Acta. 2001;312:143–51.

45. Kwon HM, Yamauchi A, Uchida S, Preston AS, Garcia-Perez A, Burg MB, et al. Cloning of the cDNa for a Na+/myo-inositol cotransporter, a hypertonicity stress protein. J Biol Chem. 1992;267:6297–301.

46. Lan X, Khandros E, Huang P, Peslak SA, Bhardwaj SK, Grevet JD, et al. The E3 ligase adaptor molecule SPOP regulates fetal hemoglobin levels in adult erythroid cells. Blood Adv. 2019;3:1586–97.

47. Li H, Handsaker B, Wysoker A, Fennell T, Ruan J, Homer N, et al. The Sequence Alignment/Map format and SAMtools. Bioinformatics. 2009;25:2078–9.

48. Li H. Minimap2: pairwise alignment for nucleotide sequences. Bioinformatics. 2018;34:3094–100.

49. Li W, Xu H, Xiao T, Cong L, Love MI, Zhang F, et al. MAGeCK enables robust identification of essential genes from genome-scale CRISPR/Cas9 knockout screens. Genome Biol. 2014;15:554.

50. Lin L, Yee SW, Kim RB, Giacomini KM. SLC transporters as therapeutic targets: emerging opportunities. Nat Rev Drug Discov. 2015;14:543–60.

51. Livermore TM, Azevedo C, Kolozsvari B, Wilson MSC, Saiardi A. Phosphate, inositol and polyphosphates. Biochem Soc Trans. 2016;44:253–9.

52. Lu B, Klingbeil O, Tarumoto Y, Somerville TDD, Huang Y-H, Wei Y, et al. A transcription factor addiction in leukemia imposed by the MLL promoter sequence. Cancer Cell. 2018;34:970–981.e8.

53. Mackay GM, Zheng L, van den Broek NJF, Gottlieb E. Analysis of Cell Metabolism Using LC-MS and Isotope Tracers. Meth Enzymol. 2015;561:171–96.

54. Martelli AM, Evangelisti C, Chiarini F, McCubrey JA. The phosphatidylinositol 3-kinase/Akt/mTOR signaling network as a therapeutic target in acute myelogenous leukemia patients. Oncotarget. 2010;1:89–103.

55. Meacham CE, Lawton LN, Soto-Feliciano YM, Pritchard JR, Joughin BA, Ehrenberger T, et al. A genome-scale in vivo loss-of-function screen identifies Phf6 as a lineage-specific regulator of leukemia cell growth. Genes Dev. 2015;29:483–8.

56. Meyers RM, Bryan JG, McFarland JM, Weir BA, Sizemore AE, Xu H, et al. Computational correction of copy number effect improves specificity of CRISPR-Cas9 essentiality screens in cancer cells. Nat Genet. 2017;49:1779–84.

57. Mulloy JC, Wunderlich M, Zheng Y, Wei J. Transforming human blood stem and progenitor cells: a new way forward in leukemia modeling. Cell Cycle. 2008;7:3314–9.

58. Nakanishi T, Balaban RS, Burg MB. Survey of osmolytes in renal cell lines. Am J Physiol. 1988;255:C181–91.

59. Nakanishi T, Tamai I. Solute carrier transporters as targets for drug delivery and pharmacological intervention for chemotherapy. J Pharm Sci. 2011;100:3731–50.

60. Nicholson LJ, Smith PR, Hiller L, Szlosarek PW, Kimberley C, Sehouli J, et al. Epigenetic silencing of argininosuccinate synthetase confers resistance to platinum-induced cell death but collateral sensitivity to arginine auxotrophy in ovarian cancer. Int J Cancer. 2009;125:1454–63

61. Nyquist MD, Prasad B, Mostaghel EA. Harnessing solute carrier transporters for precision oncology. Molecules. 2017;22.

62. Oldenborg PA, Zheleznyak A, Fang YF, Lagenaur CF, Gresham HD, Lindberg FP. Role of CD47 as a marker of self on red blood cells. Science. 2000;288:2051–4.

63. Peng H, Shen N, Qian L, Sun XL, Koduru P, Goodwin LO, et al. Hypermethylation of CpG islands in the mouse asparagine synthetase gene: relationship to asparaginase sensitivity in lymphoma cells. Partial methylation in normal cells. Br J Cancer. 2001;85:930–5.

64. Possemato R, Marks KM, Shaul YD, Pacold ME, Kim D, Birsoy K, et al. Functional genomics reveal that the serine synthesis pathway is essential in breast cancer. Nature. 2011;476:346–50.

65. Ramaley R, Fujita Y, Freese E. Purification and properties of Bacillus subtilis inositol dehydrogenase. J Biol Chem. 1979;254:7684–90.

66. Ren Y, Roy S, Ding Y, Iqbal J, Broome JD. Methylation of the asparagine synthetase promoter in human leukemic cell lines is associated with a specific methyl binding protein. Oncogene. 2004;23:3953–61.

67. Roll P, Massacrier A, Pereira S, Robaglia-Schlupp A, Cau P, Szepetowski P. New human sodium/glucose cotransporter gene (KST1): identification, characterization, and mutation analysis in ICCA (infantile convulsions and choreoathetosis) and BFIC (benign familial infantile convulsions) families. Gene. 2002;285:141–8.

68. Rossiter NJ, Huggler KS, Adelmann CH, Keys HR, Soens RW, Sabatini DM, et al. CRISPR screens in physiologic medium reveal conditionally essential genes in human cells. BioRxiv. 2020;

69. Schneider S. Inositol transport proteins. FEBS Lett. 2015;589:1049–58.

70. Shi J, Wang E, Milazzo JP, Wang Z, Kinney JB, Vakoc CR. Discovery of cancer drug targets by CRISPR-Cas9 screening of protein domains. Nat Biotechnol. 2015;33:661–7.

71. Shiue E, Prather KLJ. Improving D-glucaric acid production from myo-inositol in E. coli by increasing MIOX stability and myo-inositol transport. Metab Eng. 2014;22:22–31.

72. Sick E, Jeanne A, Schneider C, Dedieu S, Takeda K, Martiny L. CD47 update: a multifaceted actor in the tumour microenvironment of potential therapeutic interest. Br J Pharmacol. 2012;167:1415–30.

73. Simpson JT, Workman RE, Zuzarte PC, David M, Dursi LJ, Timp W. Detecting DNA cytosine methylation using nanopore sequencing. Nat Methods. 2017;14:407–10.

74. Steger DJ, Haswell ES, Miller AL, Wente SR, O’Shea EK. Regulation of chromatin remodeling by inositol polyphosphates. Science. 2003;299:114–6.

75. Stein AJ, Geiger JH. The crystal structure and mechanism of 1-L-myo-inositol-1-phosphate synthase. J Biol Chem. 2002;277:9484–91.

76. Szlosarek PW, Klabatsa A, Pallaska A, Sheaff M, Smith P, Crook T, et al. In vivo loss of expression of argininosuccinate synthetase in malignant pleural mesothelioma is a biomarker for susceptibility to arginine depletion. Clin Cancer Res. 2006;12:7126–31.

77. Tarumoto Y, Lin S, Wang J, Milazzo JP, Xu Y, Lu B, et al. Salt-inducible kinase inhibition suppresses acute myeloid leukemia progression in vivo. Blood. 2020;135:56–70.

78. Tarumoto Y, Lu B, Somerville TDD, Huang Y-H, Milazzo JP, Wu XS, et al. LKB1, Salt-Inducible Kinases, and MEF2C Are Linked Dependencies in Acute Myeloid Leukemia. Mol Cell. 2018;69:1017–1027.e6.

79. Touzart A, Lengliné E, Latiri M, Belhocine M, Smith C, Thomas X, et al. Epigenetic Silencing Affects l-Asparaginase Sensitivity and Predicts Outcome in T-ALL. Clin Cancer Res. 2019;25:2483–93.

80. Uldry M, Ibberson M, Horisberger JD, Chatton JY, Riederer BM, Thorens B. Identification of a mammalian H(+)-myo-inositol symporter expressed predominantly in the brain. EMBO J. 2001;20:4467–77.

81. Ward PS, Patel J, Wise DR, Abdel-Wahab O, Bennett BD, Coller HA, et al. The common feature of leukemia-associated IDH1 and IDH2 mutations is a neomorphic enzyme activity converting alpha-ketoglutarate to 2-hydroxyglutarate. Cancer Cell. 2010;17:225–34.

82. Wise DR, DeBerardinis RJ, Mancuso A, Sayed N, Zhang X-Y, Pfeiffer HK, et al. Myc regulates a transcriptional program that stimulates mitochondrial glutaminolysis and leads to glutamine addiction. Proc Natl Acad Sci USA. 2008;105:18782–7.

83. Wunderlich M, Mizukawa B, Chou F-S, Sexton C, Shrestha M, Saunthararajah Y, et al. AML cells are differentially sensitive to chemotherapy treatment in a human xenograft model. Blood. 2013;121:e90–7.

84. Yu W, Ye C, Greenberg ML. Inositol hexakisphosphate kinase 1 (IP6K1) regulates inositol synthesis in mammalian cells. J Biol Chem. 2016;291:10437–44.

85. Zhang Y, Zhang Y, Sun K, Meng Z, Chen L. The SLC transporter in nutrient and metabolic sensing, regulation, and drug development. J Mol Cell Biol. 2019;11:1–13.

86. Zheng S, Hou J, Zhou Y, Fang H, Wang T-T, Liu F, et al. One-pot two-strain system based on glucaric acid biosensor for rapid screening of myo-inositol oxygenase mutations and glucaric acid production in recombinant cells. Metab Eng. 2018;49:212–9.

87. Zhou L, Sheng W, Jia C, Shi X, Cao R, Wang G, et al. Musashi2 promotes the progression of pancreatic cancer through a novel ISYNA1-p21/ZEB-1 pathway. J Cell Mol Med. 2020;

